# Genomic epidemiology of *Candida auris* introduction and outbreaks in the United Kingdom

**DOI:** 10.1101/2024.01.15.575049

**Authors:** Dana Kappel, Hugh Gifford, Amelie Brackin, Alireza Abdolrasouli, David W. Eyre, Katie Jeffery, Silke Schlenz, David M. Aanensen, Colin S. Brown, Andrew Borman, Elizabeth Johnson, Alison Holmes, Darius Armstrong-James, Matthew C. Fisher, Johanna Rhodes

**Author notes:** equal contributors (Matthew C. Fisher, Johanna Rhodes).

## Abstract

**Background:** *Candida auris* is a globally emerging fungal pathogen that causes nosocomial invasive infections, particularly in intensive care units (ICU). Four prominent genetic clades originating from distinct geographic regions have been identified: South Asia (Clade I), East Asia (Clade II), Africa (Clade III) and South America (Clade IV) with each clade presenting differing antifungal resistance profiles. We aimed to elucidate the epidemiology of this infection in the United Kingdom (UK) 2014 - 2019 using genomic approaches.

**Methods:** Genome sequences from 24 isolates from six patients in four London hospitals were combined with genome sequences from 183 additional outbreak isolates from nine UK hospitals. These data were used to assess the numbers and timings of *C. auris* lineage introductions into the UK and to infer subsequent patterns of inter- and intra-hospital transmission.

**Findings:** We found evidence for at least three introductions of *C. auris* into the UK, one from Clade I and one from Clade III. The evolutionary rate of *C. auris* Clade I and Clade III were estimated at 2.764 x 10^-4^ and 3.186 x 10^-4^ substitutions per polymorphic site per year, respectively, with entry into the UK as 2013 and 2016 for Clades I and III respectively. We observed temporal and geographical evidence for multiple transmission events between hospitals and identified local within-hospital patient-to-patient transmission events.

**Interpretation:** These data confirm that *C. auris* is a newly emerged infection in the UK with at least three original introductions into this country. Our analysis shows that hospital outbreaks are linked and confirms that transmission amongst patients has occurred, explaining local hospital outbreaks. Our findings argue that enhanced surveillance of *C. auris* infection within the UK is necessary in order to protect healthcare and to curtail expansion of this emerging infection.

**Funding:** This work was supported by a Wellcome Trust Institutional Strategic Support Fund Springboard Fellowship, and by the Medical Research Council (MR/R015600/1), the Natural Environment Research Council (NE/P001165/1) and the Wellcome Trust (219551/Z/19/Z).

**Research in context:** Since its first description in 2009, *Candida auris* has spread across the globe. For this study, authors considered all publications describing whole genome sequences of isolates within the UK up until 2020 to assess the genomic epidemiology of this fungal pathogen. This study adds value to the current understanding of *C. auris* epidemiology by undertaking the first analysis to assess transmission between and within hospitals using genomic data. We also provide the first in-depth temporal analysis to estimate likely times of introduction into the UK. These results have clinical implications, encouraging hospitals to identify yeast upon admission and to assess multiple isolates from single patients, which may harbour much diversity in terms of genetics and drug resistance.

## Introduction

The ascomycete yeast *Candida auris* was first described in 2009 colonizing the ear canal of a Japanese patient (1). This fungus has now emerged as a global pathogen which has been identified in over 40 countries worldwide (2). Although often misidentified as other *Candida* species (including *C. haemulonii*), *C. auris* infections were very rare prior to 2009 (1,2) with the current global distribution explained by spatial emergence from a small number of foci. Recently, coastal waters were proposed as a potential environmental niche, with *C. auris* isolated from Andaman Islands coastal wetlands (3) and Colombian estuaries (4). *C. auris* is able to spread easily in healthcare settings, to persist for long periods, to colonise human skin, and to survive for weeks on inanimate surfaces and objects. These features of *C. auris* have led to high rates of nosocomial transmission, with outbreaks in hospitals and other healthcare facilities around the world (5–10). Whilst the intrinsic antifungal resistance of *C. auris* to fluconazole is not unusual amongst yeasts, the propensity to develop resistance to additional azole agents and also other antifungal classes make it unusual among *Candida* species (5,11,12). And, while *C. auris* mainly colonises skin, the fungus can also cause life-threatening invasive infections (13). *Candida* blood stream infections (candidaemia), are the fourth most common nosocomial bloodstream infection. For hospitalised patients with comorbidities, mortality rates exceeding 33% have been described within the first month of diagnosis with *C. auris* candidaemia, illustrating the serious nature of the infection (14). *C. auris* spread in nosocomial settings has been shown to be through contact with contaminated environmental surfaces, fomites or equipment, such as reusable temperature probes, portable ventilators and pulse oximeters, or by person-to-person contact (5,9–11,14,22). Patient movement between hospitals for rehabilitation or further treatment is thought to contribute to the spread of *C. auris* regionally and between local networks of healthcare facilities (2,28). In other countries, nationwide transmission has also been identified with genetically similar or highly identical isolates in India and Colombia being found in different hospitals, hundreds to thousands of miles apart (11,22).

Research has identified the near-simultaneous emergences of four distinct clades of *C. auris* (1). These clades are named after their inferred geographic region of origin: South Asian (Clade I), East Asian (Clade II), African (Clade III) and South American (Clade IV) with a possible fifth clade based on one isolate from Iran (15). Whilst there is low within-clade diversity, intra-clade diversity is high, with isolates between clades differing by thousands of single nucleotide polymorphisms (SNPs) (16). *C. auris* exhibits intrinsic resistance to the antifungal drug fluconazole with clade-associated point mutations in the *ERG11* gene (encoding lanosterol 14α-demethylase) associated with resistance to this drug: Y132F and K143R in Clade I, F126L in Clade III and Y132F and K143R in Clade IV (1,2,17–19). Clade II isolates, as well as the isolate from Iran (termed Clade V), are generally fluconazole-susceptible, with clade-specific patterns of azole antifungal resistance (1), varying susceptibility to amphotericin B, echinocandins and 5-flucytosine (11,18–20).

With many isolates being multidrug-resistant (MDR) and some being extensively drug-resistant (XDR), *C. auris* is a priority pathogen in the global antimicrobial resistance (AMR) agenda (21–24) and has recently been included as a priority pathogen by the WHO. Since 2015, the United Kingdom (UK) has seen numerous outbreaks in National Health Service (NHS) specialist and tertiary hospitals (5,9,25). Retrospective analysis of historical isolates at the UK National Mycology Reference Laboratory (MRL) failed to find evidence of *C. auris* in the UK prior to 2013 (26); and whilst multiple introductions of *C. auris* have been identified (27), to date the epidemiology of *C. auris* across the UK as a whole remains largely uncharacterised. At a more global scale, analysis of isolates from outbreaks and individual cases has identified countries that have isolates from different clades circulating, suggesting multiple introductions from different origins, followed by local transmission (17). In this study, we evaluate the epidemiology of *C. auris* within the UK using a genomic framework in order to determine the number and timing of introductions into the country, and to more fully understand how this pathogen transmits and spreads within and between healthcare settings.

## Methods

### Sample preparation, sequencing and data acquisition

Twenty-four isolates were collected from six patients in four London hospitals, covering an area of approx. 54 km^2^ (isolate details in minimum inhibitory concentration (MIC) antifungal susceptibility data in Supplementary Table 1) between 2016 and 2019. In addition, 43 isolates from seven centres across the UK were collected between 2013 and 2016. Isolates were confirmed as *C. auris* using matrix-assisted laser desorption/ionization-time of flight (MALD-TOF) mass spectrometry (MS). High quality, high molecular weight genomic DNA was extracted from the isolates and sent to the Sanger Institute (Wellcome Genome Campus, Cambridge, UK) for the construction of DNA libraries and sequencing. All isolates were sequenced using TruSeq Nano library preparation and Illumina HiSeq2500 sequencing of 2 x 250 bp paired end reads at The Centre for Genomic Pathogen Surveillance (Wellcome Genome Campus, Cambridge, UK). All novel raw reads in this study have been submitted to the European Nucleotide Archive under project accession PRJEB36822.

Publicly available whole genome sequences from an additional 140 isolates from across the UK were combined with these 67 newly sequenced isolates. Isolates were collected from clinical and contaminated environmental or equipment sources from nine hospitals between June 2014 and May 2019 in the UK. The samples had been sequenced using either the Illumina HiSeq 2500 platform using the HiSeq Rapid SBS Kit v2 500-cycles (Illumina) or the MiSeq platform (Illumina) using the MiSeq Reagent Kit v2 500-cycles (Illumina). All isolates included in this study are summarised in Supplementary Table 1.

### Antifungal susceptibility testing

Antifungal susceptibility testing was completed for the majority of isolates using the CLSI broth microdilution method M27-A3 (29). For a sub collection of tested isolated the susceptibility testing was carried out using Sensititre YeastOne YST-10 broth dilution panels (Thermo Fisher Scientific, UK) according to manufacturer’s instructions. The susceptibility of some isolates were determined according to the standard EUCAST method (30). MICs for isolates tested are reported in Supplementary Table 1.

### Bioinformatic analysis

Analysis was first performed for the whole UK dataset then clade-specific analyses performed separately. All isolates were initially aligned to the reference genome of B8441 from Pakistan (GenBank accession PEKT00000000.2 (1)). The data showed there were at least two introductions to the UK from two different clades, Clade I and Clade III, thus for analysis and molecular dating, the B8441 reference (GCA_002759435) was used for alignment of Clade I and reference genome B11221 (GenBank accession PGLS00000000.1 (1)) from South Africa was used for alignment of Clade III isolates using Burrows-Wheeler Aligner (BWA) v0.7.17 mem (31) and converted to sorted BAM format using SAMtools v1.16.1 (32). Duplicate reads were identified and marked using Picard v2.27.4 (33). Base Quality Score Recalibration (BQSR) was performed on all data.

Genome Analysis Toolkit (GATK) v4.2.6.1 (34) ‘HaplotypeCaller’ was used to call single nucleotide polymorphisms (SNPs) excluding repeat regions in the genome. SNPs were labelled as low confidence if they met at least one of the following parameters: DP < 5, GQ < 50, MQ < 40, MQRankSum < -12.5, ReadPosRankSum < -8.0, SOR > 4.0, or were not present in at least 90% of reads.

SNPs were converted to presence/absence data with respect to reference. Low-confidence SNPs were classed as missing. Maximum likelihood (ML) phylogenies were constructed using RAxML v8.2.9 (35) rapid bootstrap analysis over 5000 iterations using the BINCAT model of rate heterogeneity. The phylogeny of sequential isolates was constructed as above, but with 1000 bootstrap iterations using the GTRGAMMA model of rate heterogeneity. Phylogenetic trees were visualised in FigTree v1.4.4. SNPs were annotated using snpEff v5.1 (36) and summarised for *ERG11*.

### Molecular dating of *C. auris* isolates

Bayesian dating inferences and phylogenetic reconstruction were performed using BEAST v2.6.3 on whole nuclear genome alignments of 114 Clade I isolates and 93 Clade III isolates with recorded sampling dates. Due to differences in the sizes of Clade I and III *C. auris* genomes, the two clades were analysed separately. TempEst v1.5.3 (37) were used to identify significant root-to-tip distances at each node across the phylogeny. Root-to-tip regression indicating the presence of a molecular clock was identified. Therefore, to quantify the clock rates for the UK Clade I and Clade III isolates separately, Bayesian phylogenies were designed in BEAUti2 for analysis under a relaxed log normal molecular clock with a Generalised Time Reversible (GTR) nucleotide substitution model in BEAST2 v2.7.5. The specimen collection dates were used as the tip dates of the isolates and were imported as days before the present. A relaxed log normal molecular clock was chosen to account for variation in substitution rate between lineages. Bayesian Markov chain Monte Carlo (MCMC) analyses were run for 50 million steps to reach convergence in the posterior distributions (confirmed using Tracer v1.7.1) with a logging frequency of 5,000 runs and burn-in rate of 10%. Tree topology was summarised by generating a clade credibility tree in TreeAnnotator v2.6.3 with 10% burn-in and node heights set to common ancestor heights. The phylogenies were visualised in FigTree v1.4.4.

To assess the timing of entry of Clades I and III into the UK, representatives of each outbreak were assessed along with Clade I isolates B11096 and B11210 (isolated in 2014 and 2013 respectively) and Clade III isolate B11221 (isolated in 2013) (1). A HKY substitution model with optimised relaxed clock model were used, with calibrate yule model as the tree prior. A calibration node containing UK outbreak isolates (i.e. excluding B11096, B11210 and B11221) was also included. MCMC analyses were set and analysed as above.

### Outbreak transmission analysis

The R package *TransPhylo* (38) v1.4.4 uses a phylogenetic tree to infer the order and direction of transmission between isolates while taking into account within host genetic diversity and mutations common to the isolates. *TransPhylo* was used to examine transmission of isolates within each clade within the UK. MCMC analysis was run for 10,000 iterations with shape and scale parameters for the gamma distribution representing generation time and time at which observations of cases stopped. The parameters for the gamma distribution were set to 10 and 1, respectively. A generation time of 10 days was assumed to be plausible in an inpatient setting. Convergence was assessed by plotting the tree output and the “CODA” package (39) in R version 4.0.4. Transmission trees and probability of direct transmission for all pairs of individuals were plotted in R.

## Results

Two hundred and seven isolates collected from nine hospitals between June 2014 and May 2019 in the UK, including the 67 newly sequenced isolates, were aligned to the *C. auris* reference genome, B8441 (Clade I). Out of the 207 isolates, 114 (55%) were from Clade I and 93 (45%) were from Clade III. Of the sequenced isolates, 187 were isolated from patients, 13 from hospital environments and 7 were of unknown origin. In two hospitals (King’s College Hospital and Wexham Park Hospital), isolates from both clades were co-circulating. Clade III isolates were subsequently realigned to the B11221 (Clade III) reference genome. The mean genome coverage was 61x (range 15x to 227x) for Clade I and 45x (range 7x to 118x) for Clade III. On average, for Clade I, 99.14% and for Clade III, 98.22% of all reads were successfully mapped to their respective reference genomes. After two filtering steps to retain only high confidence SNPs, on average for Clade I and Clade III, 94.2% and 92.9% SNPs were kept.

Nonsynonymous substitutions within the coding regions of the *ERG11*, *FKS1* and *FUR1* genes were evaluated to search for known mutations conferring resistance to fluconazole, echinocandins and flucytosine antifungal drugs, respectively. Fluconazole resistance associated mutations in the *ERG11* region were found in all isolates (Figure 1); 33% (*n* = 69) of isolates have been MIC tested for fluconazole (Supplementary Table 1), and all displayed fluconazole resistance. Of the 207 isolates, 44.9% (*n* = 93) had the F126L *ERG11* substitution, of which all isolates were in Clade III; while 6.3% (*n* = 13) had the K143R *ERG11* substitution, and 48.8% (*n* = 101) had the Y132F *ERG11* substitution, and were found in Clade I (Table 1). The point mutations in the *ERG11* gene in the UK are clade-specific with Clade I exhibiting K143R and Y132F and Clade III exhibiting F126L substitutions. This distinction does not expand globally as Clade IV has the same resistance mutations in *ERG11* as Clade I (1,16,17). All Clade III isolates also contained the Y125A substitution within *ERG11*. This allele has not been found previously in *C. auris*. Analysis of the *FKS1* and *FUR1* genes showed no mutations. However, 13 isolates displayed elevated MICs for flucytosine, which may be indicative of drug resistance, suggesting alternate mechanisms other than *FUR1* being responsible. None of the isolates with completed antifungal susceptibility testing displayed raised MICs to echinocandin antifungal drugs. A summary of the data can be found in Supplementary Table 1.

**Figure 1.**
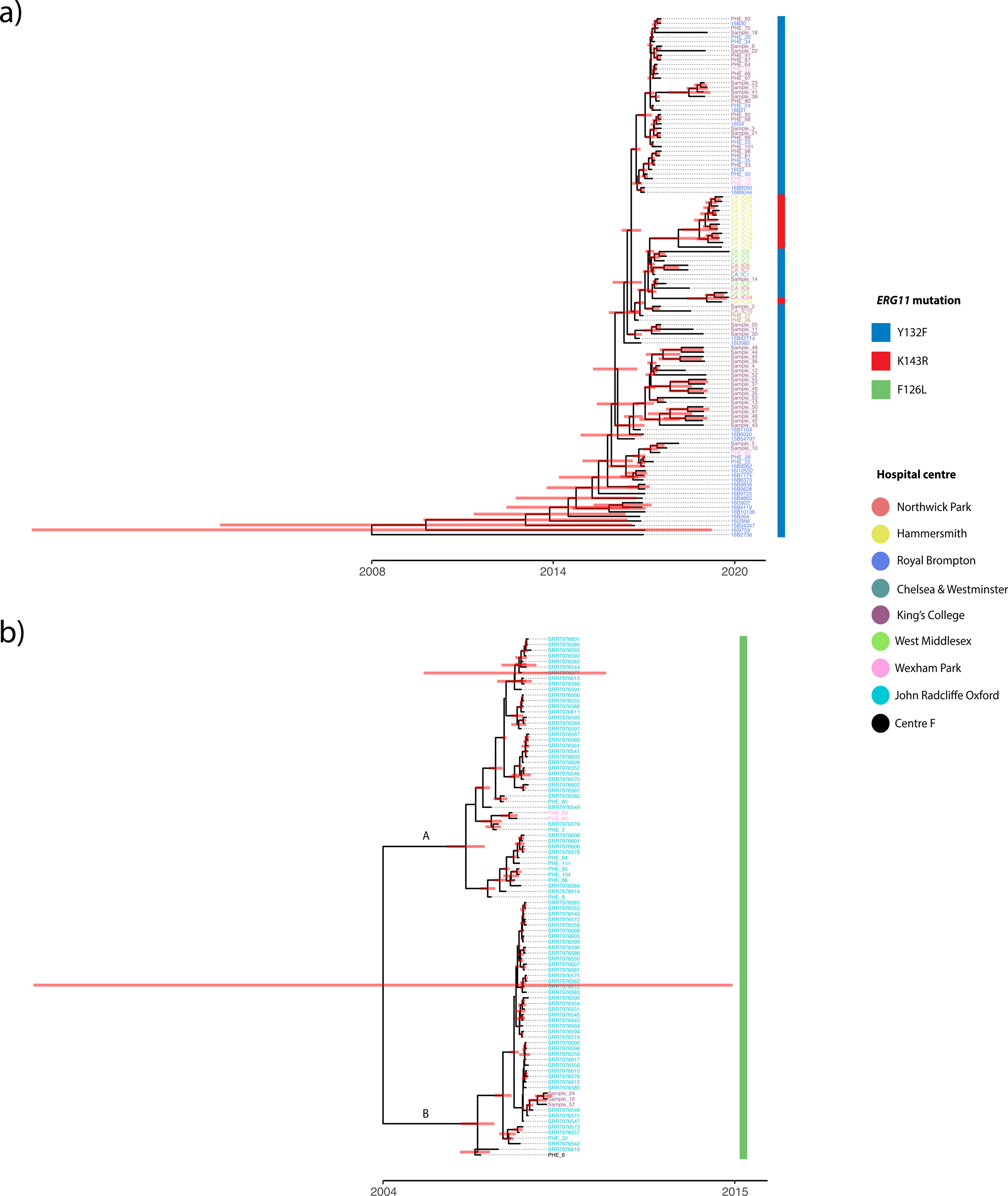
Dating the introduction of *C. auris* into the United Kingdom. *ERG11* mutations are shown for each isolate **a)** Maximum Clade Credibility phylogeny using the posterior tree distributions of UK Clade I isolates showing a TMRCA as 2008 (CI: 1989-2015) **b)** Maximum Clade Credibility phylogeny using the posterior tree distributions of UK Clade III isolates showing a TMRCA as 2004 (CI: 1972-2015). Subclades A and B are denoted and posterior distributions shown.

**Table 1.**
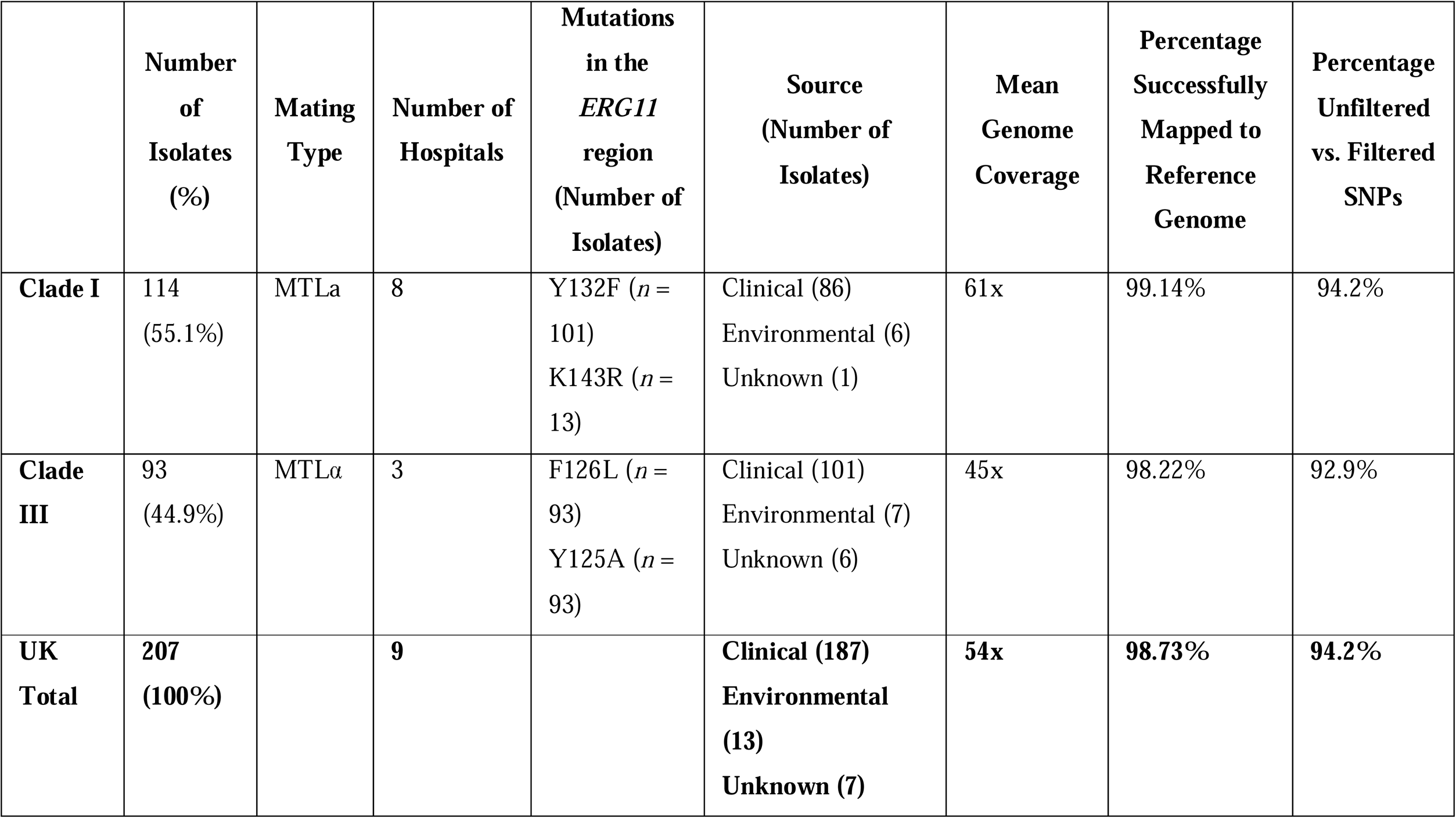
Summary of Clade I and Clade III isolates included within this study and details on *ERG11* polymorphisms.

### Temporal analysis shows at least three introductions of *C. auris* into the United Kingdom

Maximum likelihood (ML) trees inferred using whole-genome SNPs for Clade I and III were used as the input to calculate root-to-tip regressions for each clade (Supplementary Figure 1). The analysis included all 114 Clade I and 93 Clade III isolates from the UK dataset, with the date of sample collection used for tip dating. We observed a positive correlation between the tip dates and divergence from the root using TempEst for Clade I (R^2^ = 0.38, *p*-value 1.93e^-13^), and Clade III (R^2^ = 0.53, *p*-value 6.19e^-17^). Phylostems estimated the TMRCA corresponding to the x-intercept of the regression, indicating that the MRCA for Clade I and Clade III occurred in 2000 and 2013 respectively, representing the most likely year of entry into the UK if each clade was only introduced on a single occasion.

To refine this estimate, the dataset was subjected to coalescence analysis using BEAST. Using the posterior distributions of the Clade I and III phylogenies, Maximum Credibility Clade trees (MCC) were inferred to analyse introductions into the UK and TMRCA (Figure 1).

For Clade I, the strong temporal signal, basal position of the first hospital outbreak isolate (2015) and ladderised MCC phylogeny suggests the TMRCA of this clade in the UK as 2008 (95% highest posterior density (HPD) interval [1989, 2015]) followed by its spread. The clock rate was defined as 2.764x10^-04^. Assessing the introduction of Clade I into the UK placed entry mid-September 2013 (95% HPD late November 2012 to mid-September 2014 with a posterior probability (PP) of 1). Interestingly, the two resistance-associated *ERG11* point mutations (K143R and Y132F) do not cluster in the phylogeny suggesting that these SNPs may have originated more than once. Based on the Clade I MCC tree, introduction and diversification in the UK began at Royal Brompton Hospital and spread to the other hospitals, including seeding the later Hammersmith Hospital outbreak that includes the K143R *ERG11* allele. Of note, the branch leading to the Hammersmith Hospital cluster and containing the K143R mutation has a high posterior probability (PP=0.9202) indicating an introduction to the hospital in 2018 (95% HPD [2017, 2019]); this compares well with the detection of this outbreak in November 2018. Similarly, a group of isolates from one of the two inferred introductions into King’s College Hospital is also very well supported (PP=0.9892) with the TMRCA approximately in 2016 (95% HPD [2016, 2017] comparing well with the detection of this outbreak in April 2016 (25).

The topology of the Clade III tree (Figure 1b) suggests at least two introductions into the UK due to the bifurcating phylogeny and low-quality temporal signal. The long basal branch lengths of the Clade III phylogeny indicate that if a single introduction in 2004 had occurred, it would have remained undetected for approximately 2-3 years, which is clinically unlikely. The good posterior support of the two sub-clades (Figure 1b) in late 2006 suggests two introductions of Clade III near-simultaneously, into the UK remains more likely. The clock rate was defined as 3.186x10^-04^. Assessing the time-scaled phylogeny placed entry into the UK late 2014 (95% HPD early Oct 2014 to Nov 2014 with a posterior probability of 1); the earliest Clade III UK isolate (PHE_6) was isolated in 2014, making these dates likely. Only an average of 379 SNPs separate the two Clade III subclades, yet the SNP distribution for each subclade differs; subclade-specific missense variants mapped to 33 and 51 genes unique to subclade A and B respectively. Whilst no significant corrected gene ontology (GO) terms were associated with the genes in subclade B, the genes in subclade A were significantly overrepresented for the biological process cell wall (1 ➔ 3)-β-D-glucan biosynthetic process. These differences are consistent with two initial separate introductions of Clade III into the UK, with both subclades introduced into John Radcliffe hospital and diversification and possible spread to King’s College and Wrexham Park hospitals.

### Inferring direct and indirect patient-to-patient transmission of *C. auris*

Inter- and intra-hospital transmission was analysed using the R package *TransPhylo*. A matrix of probability of direct transmission for all pairs of individual isolates within each clade were generated to evaluate transmission events within and between hospital centres (Supplementary Figure 3). We detected 28 (Clade I) and 82 (Clade III) events with a transmission probability of greater than 75%. Clade I events included movements between seven of the eight hospital centres, however two of the transmission events inferred occurred prior to detection indicating either false implied directionality or sampling did not accurately elucidate temporal trends in these data. Clade III transmission events included all three hospital centres; however, 33 of the 82 events were inferred to have occurred prior to the collection date of the baseline isolate.

Sixteen transmission events occurred within Clade I amongst patients within hospitals while 12 transmission events occurred between hospitals (Figure 2a, Supplementary Table 3). Twenty-three transmission events in Clade I were robustly supported with a transmission probability of 100%, including 13 within hospital and 10 between hospital events; two events also showed links between clinical and environmental isolates (e.g. inanimate surfaces), underscoring the environmental risk that *C. auris* presents in hospital settings. Of the 13 within hospital transmission events, 12 were transmission between patients and one was transmission between a patient and an environmental surface. Based on the patterns of the 12 between hospital events, isolates originated in Royal Brompton Hospital before being transmitted to King’s College and Chelsea and Westminster hospitals then later to Wexham Park and Northwick Park hospitals across London. Of note, the Hammersmith Hospital cluster with the K143R *ERG11* mutation appears to be self-contained and was only found circulating within this hospital.

**Figure 2.**
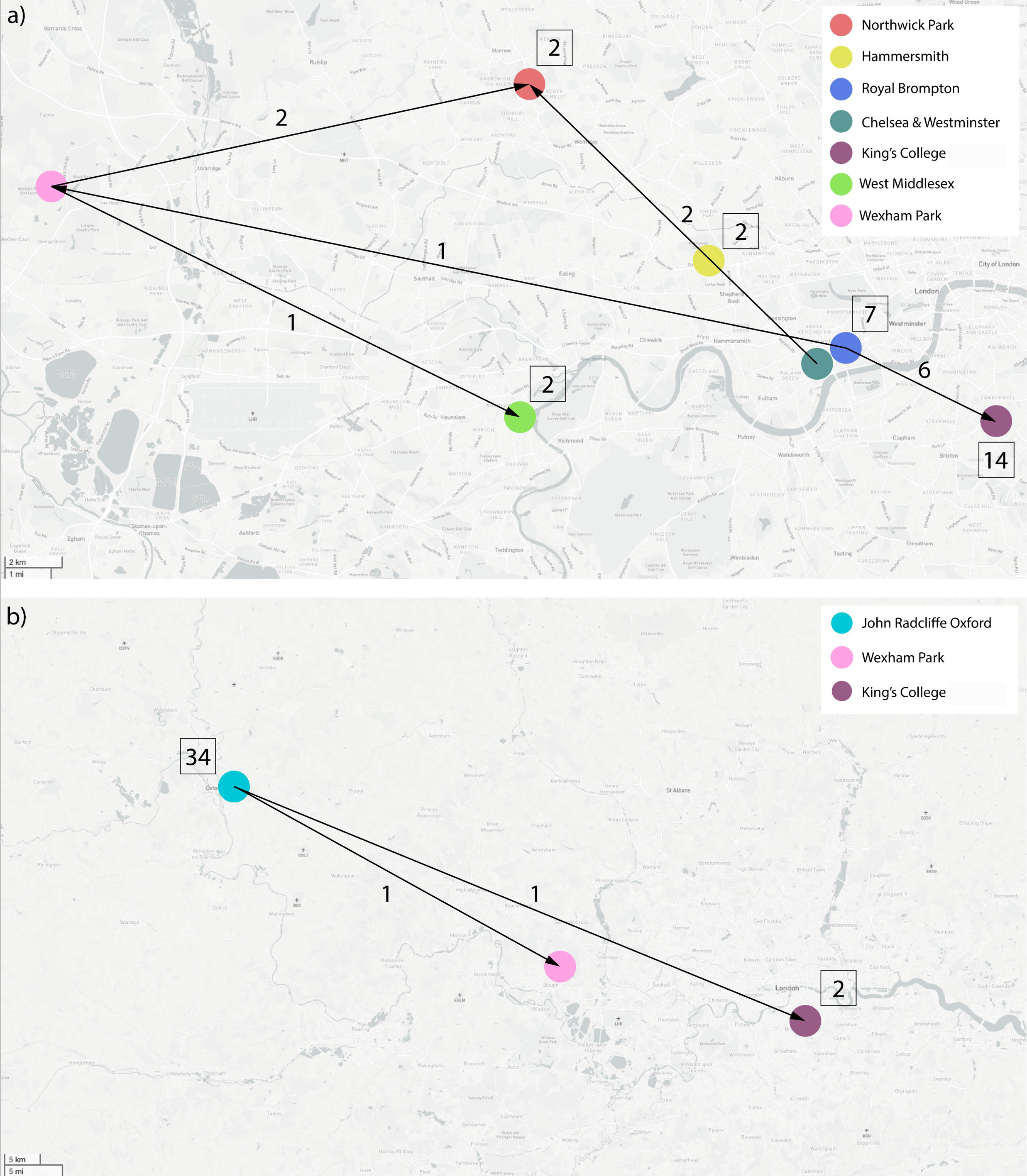
Intra- and Inter-hospital transmission events identified by TransPhylo. **a)** Clade I direct transmission directionality according to the analysis in TransPhylo of within hospitals between patients or environmental surfaces and between hospitals **b)** Clade III direct transmission directionality according to the analysis in TransPhylo of within hospitals between patients or environmental surfaces and between hospitals. In both, numbers above arrows indicate number of identified inter-hospital transmissions, whilst numbers within squares indicate number of identified intra-hospital transmissions.

Eighty transmission events within Clade III occurred within hospitals while two transmission events occurred between hospitals (Figure 2b, Supplementary Table 4). Sixty transmission events in Clade III had a transmission probability of 100%, including 59 within hospital and one between hospital event; three events also showed movement between clinical and environmental isolates. Of these transmission events, five were multiple isolates within the same patient taken from different sources on different dates. Additional transmission events were between patients and five were transmission events between a patient and an environmental surface. Based on the two intra-hospital transmission events, transmission started in John Radcliffe Hospital and moved to King’s College and Wexham Park hospitals, which is consistent with the findings of the temporal analysis.

### Microevolution of *C. auris* within a single patient

Twenty-four *C. auris* isolates sequenced as part of this study were taken sequentially from six patients admitted to four London hospitals to investigate microevolution within the human host (Supplementary Table 5). All isolates displayed raised MICs to fluconazole (Supplementary Table 1).

Five isolates were sequenced from the same patient in West Middlesex Hospital over a period of 29 months (December 2016 – May 2019) at multiple bodily locations. Isolates were separated by an average of 8 SNPs, whilst phylogenetic analysis suggests two distinct clusters based on body location with substantial bootstrap support (Supplementary Figure 4): nose (CA_IC3 and 4), and urine (CA_IC2, 5 and 6). Nose and urine isolates were separated by 3 and 7 SNPs respectively. Analysing nose and urine isolates separately, only CAIC6 had accumulated non-synonymous SNPs (nsSNPs) when compared to the initial urinary isolate (CA_IC2), with 2 nsSNPs in B9J08_003363 (Figure 3a), and ncRNA gene tRNA binding lysine. Only intergenic SNPs were found to be unique to either of the two nose isolates (Figure 3a, Case 2).

**Figure 3.**
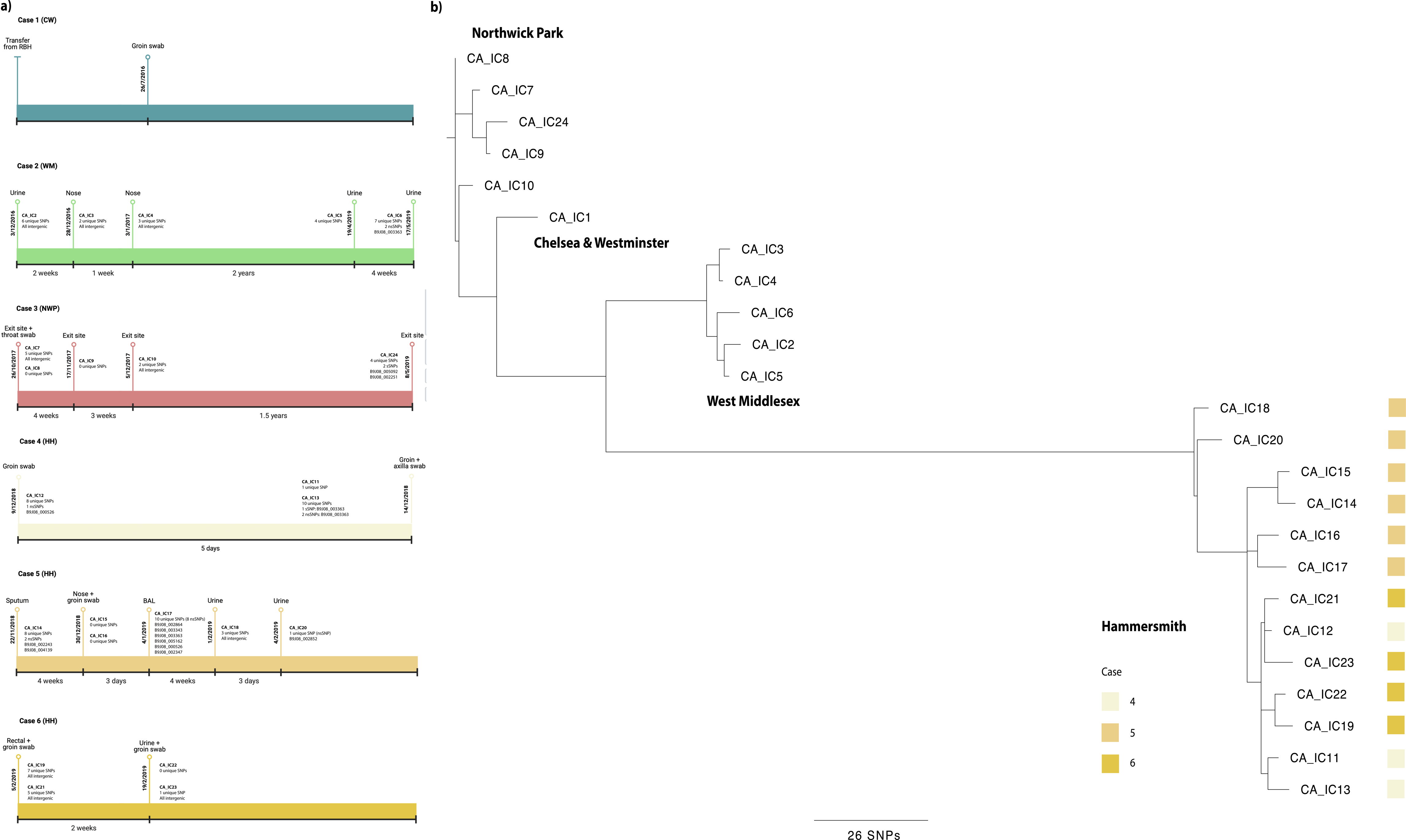
Microevolution of sequential isolates. **a)** timelines of six patients in four London hospitals with sampling dates, location of sample and unique SNPs accumulated. **b)** unrooted, ordered, maximum likelihood phylogenetic tree constructed in RAxML using whole-genome SNPs for sequential isolates taken from six patients in four London hospitals. Branch lengths represent the average number of SNPs.

Five isolates, from a single throat swab and catheter exit sites, were sampled from a single patient at Northwick Park Hospital over a period of 18 months (October 2017 – May 2019). Despite being collected 18 months after the original four isolates (CA_IC7-10), CA_IC24 is more closely related to CA_IC7-9, separated by only 7 SNPs on average, whilst CA_IC10 is 8 SNPs on average separated from isolates CA_IC7-9 and 24. CA_IC10 potentially represents infection from a genetically distinct *C. auris* variant; transmission analysis via *TransPhylo* corroborates this, identifying a significant intra-hospital transmission of isolate Sample_2 from King’s College Hospital to Northwick Park Hospital (Supplementary Table 4). Isolates CA_IC7-10 are also identified as significant transmission events, suggesting primary colonization by a single isolate subsequently colonising the patient. These isolates were compared to the initial throat and exit site swabs taken in October 2017, and only CA_IC24 had unique synonymous SNPs, which mapped to two hypothetical genes, B9J08_002251 and B9J08_005092 (Figure 3a, Case 3).

Sequential isolates were taken from three other patients (Cases 4-6, Figure 3a) at Hammersmith hospital (HH) over four months. Phylogenetic analysis shows some dissimilarities between isolates from the same patient over time (Figure 3b): in case 4, the two samples taken on 14^th^ December 2018 (axilla and groin swabs, CA_IC11 and CA_IC13 respectively) are genetically closely related, separated by only 7 SNPs, yet distinct from the initial groin swab (CA_IC12) taken five days prior. Indeed, CA_IC12 is only separated by 7 SNPs from CA_IC21 and 23 from Case 6. However, given the high number of unique SNPs in each isolate (Figure 3a), it is probable this patient was infected with at least three genotypes. Similar to Case 2 (WM), unique synonymous and nsSNPs were observed in the ncRNA tRNA-Lys gene B9J08_003363 in groin isolate CA_IC13. The genetically dissimilar isolates from Case 5, six taken from multiple anatomical sites over 3 months, also seems to indicate colonisation by a diverse *C. auris* genetic population: on average isolates were separated by 17 SNPs, yet the two urine isolates (CA_IC18 and CA_IC20, collected one month apart) are only 9 SNPs different, but 23 SNPs apart from other Case 5 isolates.

Temporal analysis (Figure 1) also places these two isolates as basal for HH, suggesting that whilst these two isolates were taken last, they potentially represent the original infecting population. More unique SNPs were recovered in this patient, with B9J08_003363 again displaying unique nsSNPs in CA_IC17, isolated from a bronchoalveolar lavage (BAL) fluid sample. Finally, four isolates over a two-week time period were collected from Case 6 at HH and were on average 9 SNPs apart. The first 3 isolates (CA_IC19, 21, 22) were groin and rectal swabs, whereas the final isolate CA_IC23 was from a urine sample. Comparison between early and later isolates yielded only unique SNPs mapping to intergenic regions (Figure 3a).

## Discussion

Our study shows that there have been multiple introductions from different spatial origins at different times into the UK followed by within and between-hospital transmission. Only Clades I and III, the South Asian and African clades respectively, were recovered within the UK based on phylogenetic analysis and clade specific *ERG11* gene mutations. Using the whole-genome SNPs, Bayesian inference determined the MRCA of the two clades to be in 2008 for Clade I and 2004 for Clade III. Introduction into the UK was placed at mid-September 2013 and late 2014 for Clade I and III, respectively. Local-scale transmission patterns within and between hospitals were identified in both clades showing transmission of *C. auris* from human-to-human and between human and environmental surfaces. Analysis of the microevolution of sequentially collected isolates from single patients in multiple hospitals has shown patients can be colonised and/or infected by a diverse genotype, confirming earlier studies (19). It is therefore important to collect and sequence multiple *C. auris* per patient, ideally from different body sites, to fully assess the genetic diversity present; whilst difference in drug susceptibility profiles were not observed in sequential isolates in this study, it cannot be ruled out as a possibility, which could negatively impact patient treatment. Interestingly, nsSNPs within B9J08_003363, encoded a ncRNA tRNA-Lys, were identified in multiple isolates from different patients and hospitals (CA_IC6, WM; CA_IC13, Case 4 HH; CA_IC17, Case 5 HH). ncRNAs have been shown to regulate gene expression in response to various conditions, such as antifungal drugs, in what is termed as ‘epimutation’ (40,41). This finding could be explored further to investigate the role of ncRNAs in *C. auris* adaptation within the human host.

The slope of the root-to-tip regression of Clade I and III were found to be 4.901x10^-4^ and 3.186x10^-4^ substitutions per polymorphic site per year, respectively. These evolutionary rates are reasonably comparable to other epidemiological analyses of *C. auris*, the Royal Brompton outbreak (1.002x10^-3^ substitutions per polymorphic site per year) and a global dataset including 304 isolates (1.8695x10^-5^ substitutions per polymorphic site per year) (17,19). The Royal Brompton Hospital outbreak included the 24 outbreak-specific Clade I isolates and used a relaxed lognormal molecular clock, which accounts for variation in evolution, while the analysis of the global dataset used isolates from all four clades using a strict molecular clock, which assumes a constant rate of evolution. These differences may account for the variability between the two evolutionary rates predicted for *C. auris* in previous studies. A relaxed lognormal molecular clock was also used in this study to account for variation in substitution rates, which has been suggested as an appropriate clock in fungi (42).

While support for two clades in the UK is well-documented based on hospital-specific outbreak analysis, this is the first-time temporal analysis on all available Clade I and III WGS data in the UK has been performed. According to the Bayesian MCC tree, there was a single introduction with a MRCA in Clade I estimated to be in 2008 (95% HPD interval [1989, 2015]). The Bayesian phylogenetic analysis shows that the first introduction of Clade I into the UK was at RBH with multiple introductions into the hospital consistent with the conclusions from our original study (19), due to the scattered isolates throughout the time-scaled phylogeny. This is consistent with the RBH outbreak analysis which suggested that entry into the hospital was in early 2015 before all other hospitals with Clade I isolates (19). However, the long branches leading to the isolate on either side of the root have low posterior probabilities (PP=0.2044) showing poor support for that topology, suggesting these isolates could have been circulating within the hospital before detection. This observation highlights the importance of yeast identification in clinical samples. The introduction of Clade I into the UK does not differentiate between the isolates with the K143R and Y132F point mutations in the *ERG11* gene, despite isolates from Hammersmith Hospital containing the K143R mutation has a very well-supported branch. The MRCA for this hospital is estimated in late 2018 (95% HPD [2017, 2019]), which corresponds to the sample collection dates in the hospital (Supplementary Table 1). Transmission analysis with *TransPhylo* also suggested the Hammersmith Hospital cluster was self-contained. However, there is not sufficient evidence here to confirm this genotype represents a separate introduction into the UK.

The MRCA in Clade III was estimated to be in 2004 (95% HPD interval [1972, 2015]); however, given the bi-clade structure with long basal branches, it is more indicative that the diversity of Clade III has not been fully sampled here. Clade III was initially introduced into the John Radcliffe hospital via two introductions before spreading to King’s College Hospital and Wrexham Park Hospital. This is at odds with the original analysis of Eyre *et al.* which concluded the outbreak originated from a single introduction from Clade III in 2013 (5). This may be due to differing clock rates. Since the John Radcliffe hospital only serves 1% of the UK population, two introductions may be likely if a single patient brought in pre-existing diversity, or colonisation went undetected. However, the distribution of non-synonymous SNPs in Clade III sub-clade A, which was introduced into the UK slightly earlier than sub-clade B, is significantly overrepresented for the β-D-glucan biosynthetic process, a crucial component of the cell wall. Non-synonymous mutations in genes involved in β-D-glucan biosynthesis could represent evolution for these isolates to have higher β-D-glucan content in their cell walls. *Aspergillus flavus* isolates with higher β-D-glucan content may show amphotericin B resistance (43); further analysis into cell wall content needs to be carried out to confirm difference in Clade III isolates within the UK.

As nosocomial transmission of *C. auris* has been highlighted as such an important factor in healthcare-associated infection control measures, reconstructing significant transmission events from genomic data can be informative. We applied *TransPhylo* (44), a software tool implemented as an R package, to link the *C. auris* phylogenies and predicted transmission events, and identified 28 and 82 individual transmission events for Clade I and III, respectively, based on the 75% probability threshold of direct transmission of isolates (Figure 2). Since *TransPhylo* was intended to be applied to bacterial pathogens, which can have clonal genomes, *C. auris* was deemed to be a logical dataset for application of this =software. In Clade I, 16 of the events occurred within hospitals while 12 occurred between hospitals (Supplementary Table 3). Between-hospital patterns of transmission show that the first observed occurrence of Clade I in the UK started in Royal Brompton Hospital, confirming that this is the European index outbreak for *C. auris*. The transmission analysis was also able to identify the HH isolates with the K143R point mutation in the *ERG11* gene as a separate transmission chain circulating only within the hospital. TransPhylo’s built-in algorithm to identify introductions into a population of interest (38) allowed the identification of the separate transmission chain and unsampled intermediates for the different Clade I *ERG11* point mutations. By comparing the transmission events in Clade III to the corresponding MCC phylogeny, we showed that all transmission events stayed within their corresponding location on either side of the MCC root (Supplementary Table 4). Eighty of the 82 events occurred within hospitals, with four genotypes accounting for 79.3% of all transmission events. Two inter-hospital transmissions showed patient/isolate movement from John Radcliffe to both KCH and WPH. Analysis from the John Radcliffe outbreak confirmed that the isolates from temperature probes (environmental surface) were scattered throughout the phylogeny, and there was no evidence of transmission from patient-to-patient in nearby beds (5); here, only four of the 82 transmission events were shown to occur between the temperature probes and patients, although this is likely a result of under sampling of temperature probes early in the outbreak. Earlier analyses of this outbreak using WGS failed to find a relationship between patient bed proximity and genomic distances, which may be an artefact of having only a few environmental samples obtained later in the outbreak.

A common limitation that confounds inference of transmission events here is that the sample collection date of the second isolate occurs before the collection date of the primary isolate. There are multiple reasons why this could transpire. It is possible that the sample size for each of the clades was not large enough to evaluate due to missing intermediate isolates on the pathway to transmission. The movement of hospital equipment, staff and patients contaminated with *C. auris* from hospital to hospital could affect the sample collection dates and transmission between hospitals; for example, if a patient with chronic undetected asymptomatic *C. auris* contaminated a piece of equipment that was then used on an independent patient who then developed symptomatic infection, the sample collection date for the second patient in this example would be prior to the first patient who would not be identified with *C. auris* until after they show symptoms, are detected by contact tracing or possibly never. Also, the amount of community carriage is difficult, or impossible, to assess. Finally, *C. auris* is incredibly resilient, often showing resistance to disinfectants and can form biofilms allowing it to remain dormant and persist on surfaces for long durations possibly allowing patients to be infected by the same isolate months apart, although this could also result from undetected chains of transmission (2,5,16,19).

The present analysis evaluated the spatial and temporal patterns of Clade I and III isolates of *C. auris* within the UK. Future analyses would benefit from comparing these isolates with the full global dataset to help expand understanding of where the UK fits within the global phylogenetic tree. By including within-clade introductions and differentiation between the *ERG11* mutations we could also understand the temporal origins of outbreaks and infections with Y132F and K143R mutations in the *ERG11* gene. In addition, the use of patient, equipment and staff records, after ethics approval is granted, to validate the outputs from TransPhylo can be used to demonstrate the capability of identifying transmission patterns retrospectively. If accurate, TransPhylo can also be utilised to predict local-transmission dynamics prospectively in the beginning of outbreaks to help in the earlier implementation of infection prevention and control measures and containment of the outbreak.

This study is the first to assess the introduction of *C. auris* into the UK, which has experienced outbreaks since 2015 of isolates from two of the five clades (Clades I and III) (26), and a single introduction of Clade II, using whole genome sequencing. Leveraging the power of whole genome sequencing we identified at least three introductions into the UK, and importantly, provide novel insights into inter- and intra-hospital transmission of *C. auris*. These analyses underscore the need to utilise the vast numbers of genome sequences available for *C. auris* to further assess transmission routes into and between hospitals and help to inform hospital policy on outbreak management.

## Supporting information

SuppInfo

---

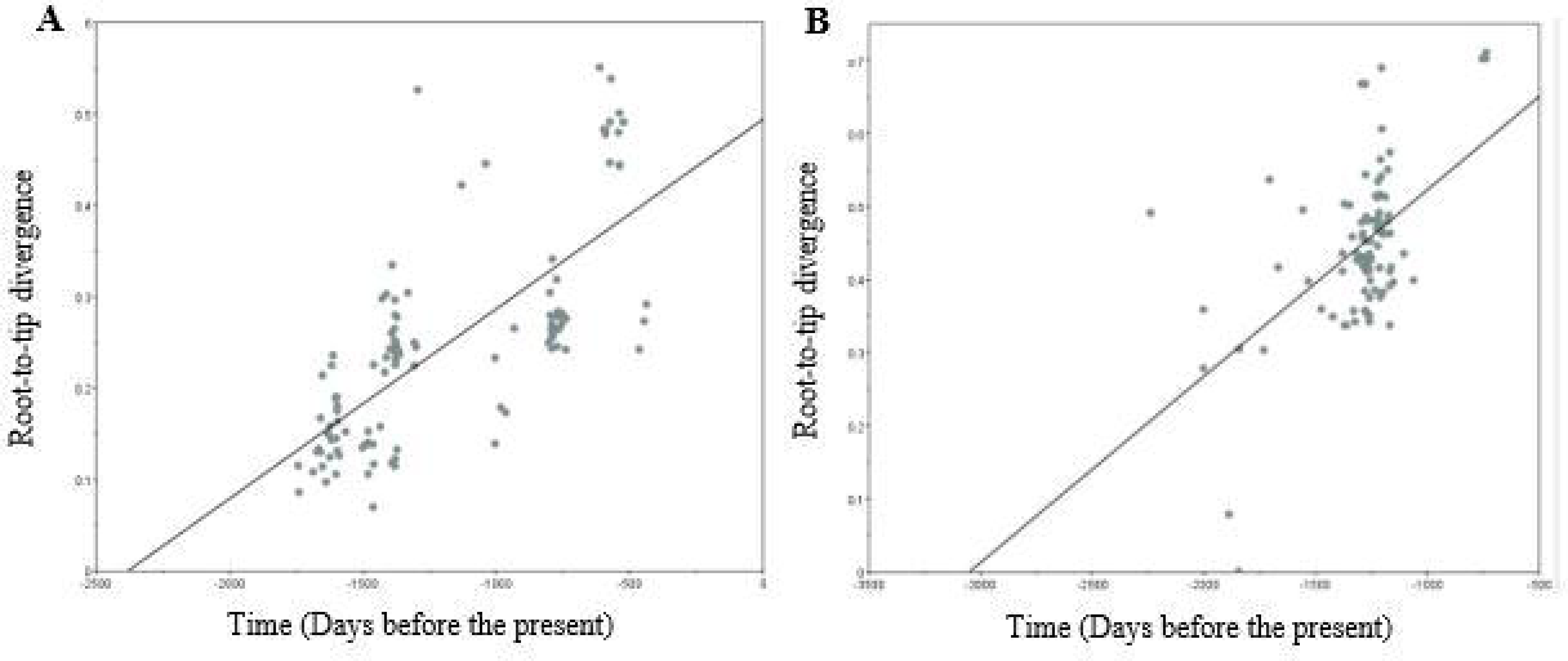

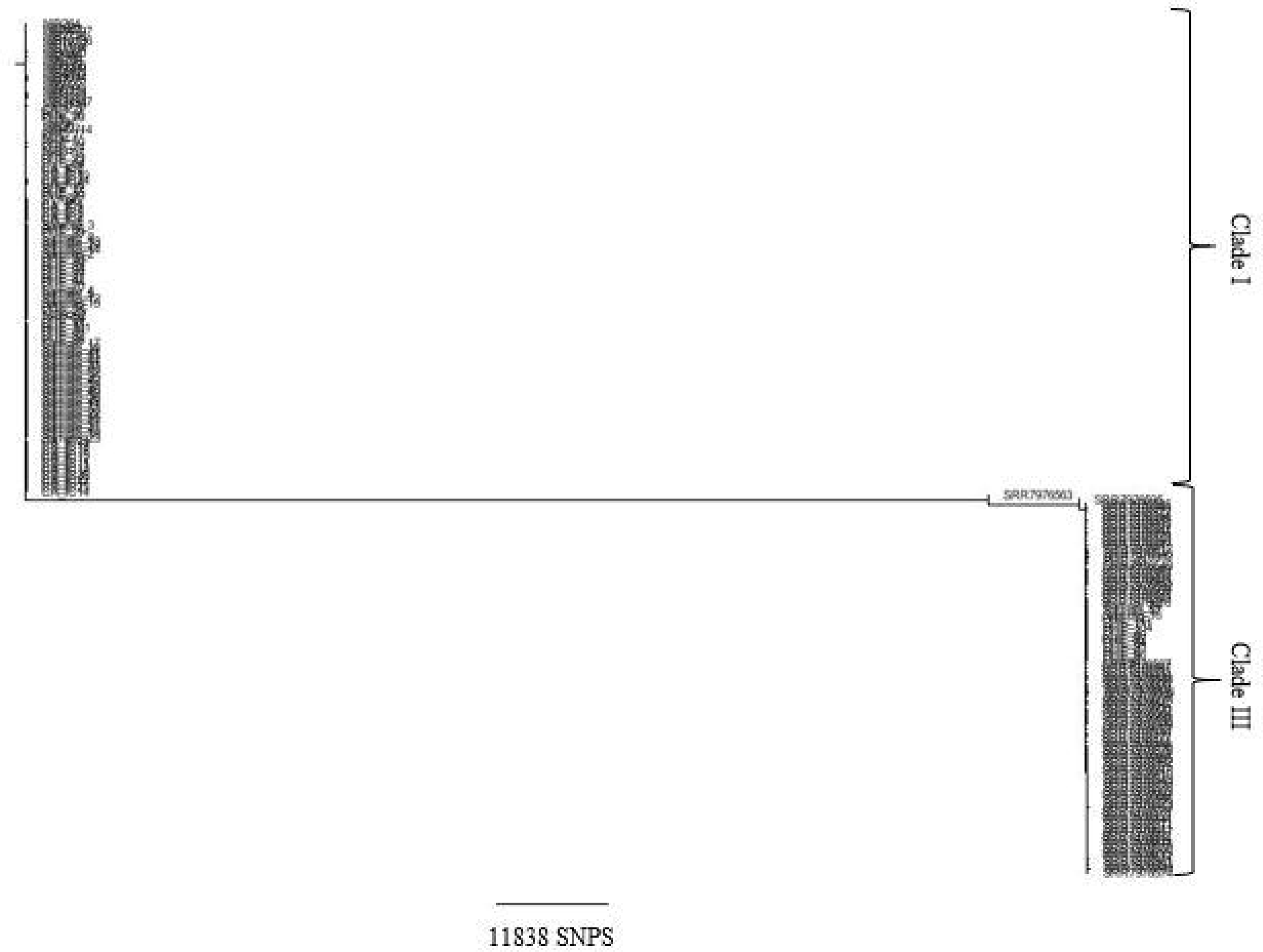

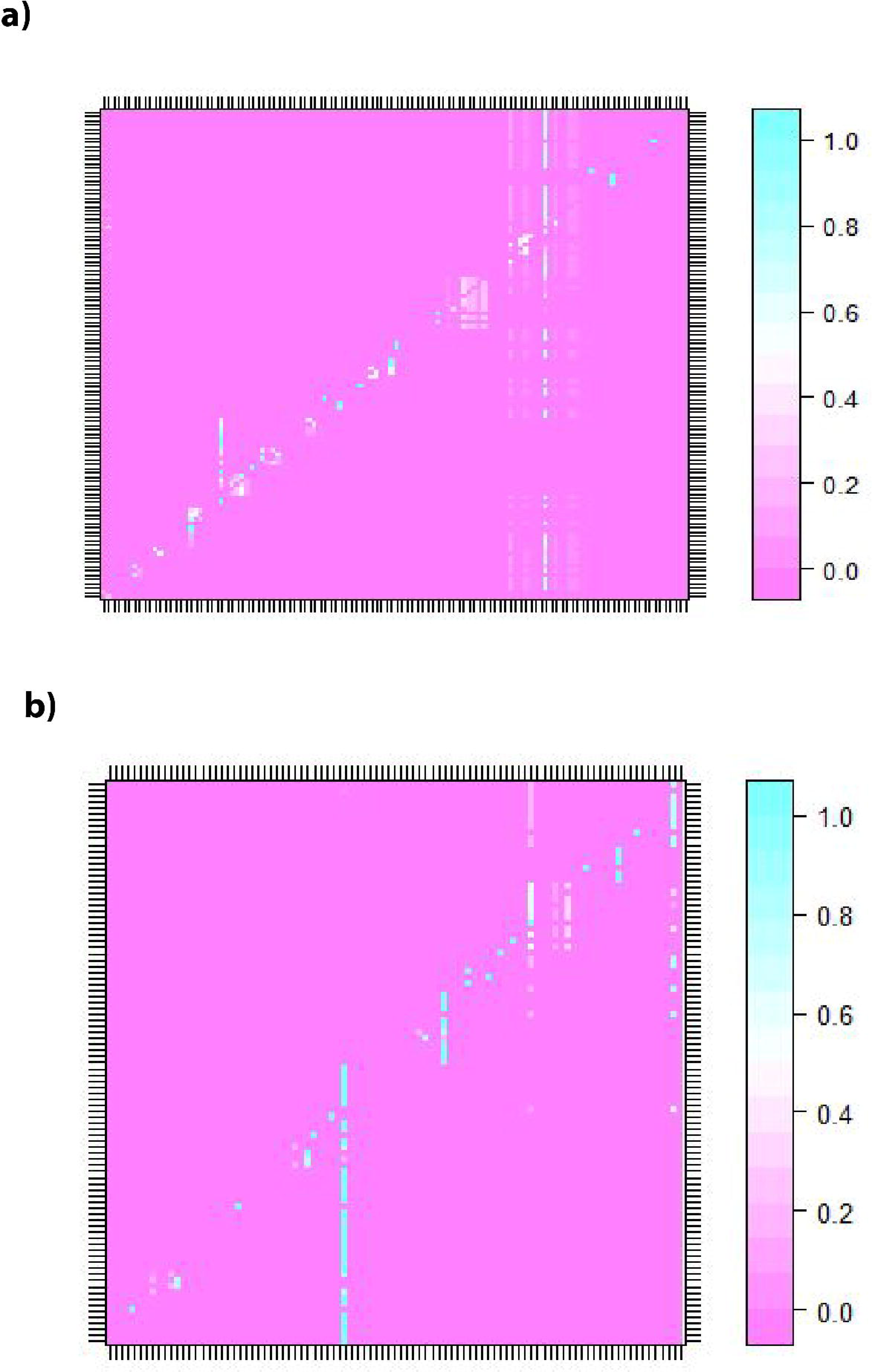

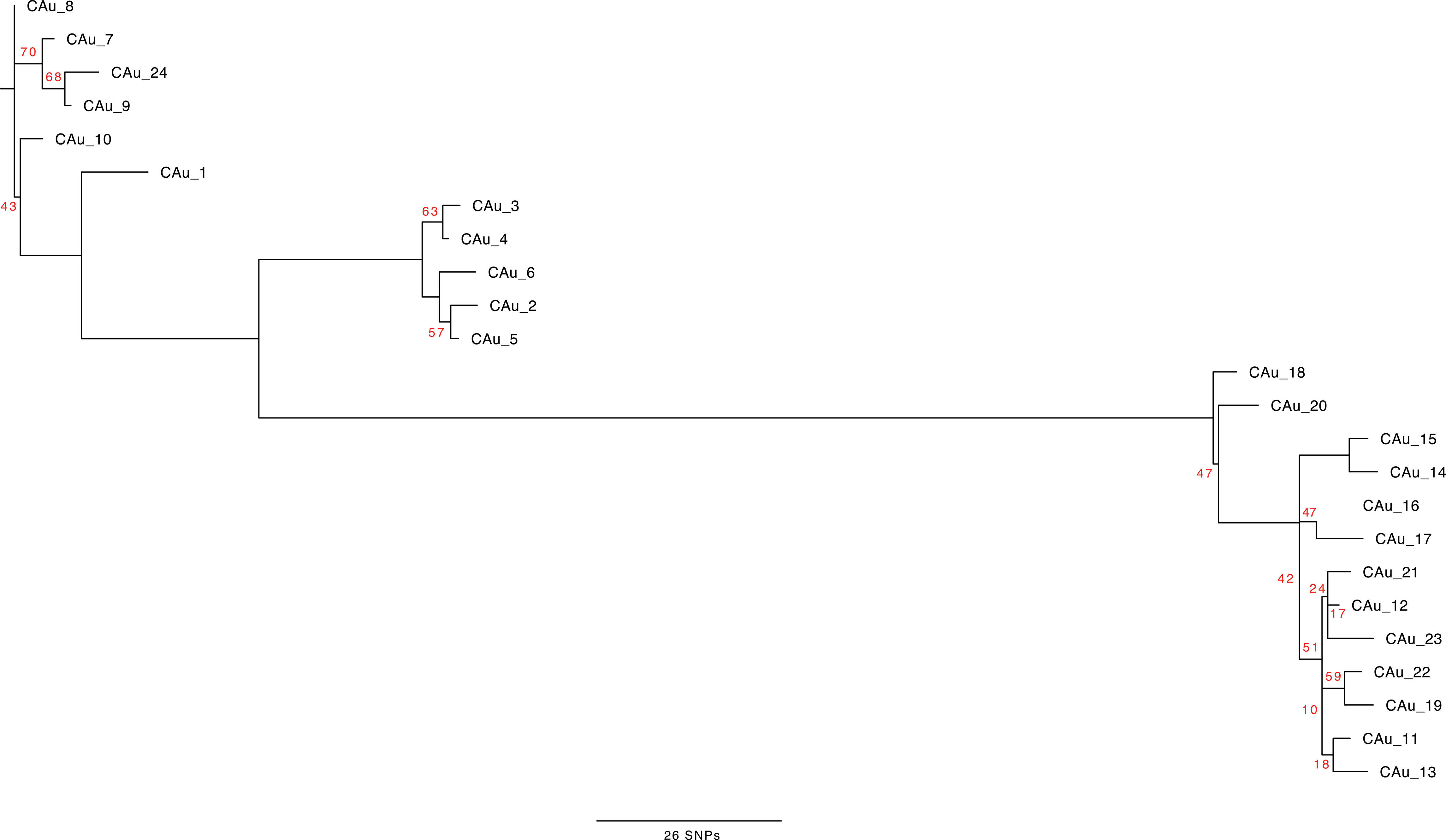

## References

1. Lockhart SR, Etienne KA, Vallabhaneni S, Farooqi J, Chowdhary A, Govender NP, et al. Simultaneous emergence of multidrug resistant Candida auris on three continents confirmed by whole genome sequencing and epidemiological analyses. Clin Infect Dis. 2017;64(2):134–40.

2. Rhodes J, Fisher MC. Global epidemiology of emerging Candida auris. Curr Opin Microbiol. 2019 Dec 1;52:84–9.

3. Arora P, Singh P, Wang Y, Yadav A, Pawar K, Singh A, et al. Environmental Isolation of Candida auris from the Coastal Wetlands of Andaman Islands, India. mBio. 2021;12(2):e03181–20.

4. Escandón P. Novel Environmental Niches for Candida auris: Isolation from a Coastal Habitat in Colombia. J Fungi. 2022 Jul;8(7):748.

5. Eyre DW, Sheppard AE, Madder H, Moir I, Moroney R, Quan TP, et al. A Candida auris Outbreak and Its Control in an Intensive Care Setting. N Engl J Med. 2018;379(14):1322– 31.

6. Calvo B, Melo AS de A, Perozo-Mena A, Hernandez M, Francisco EC, Hagen F, et al. First report of Candida auris in America: clinical and microbiological aspects of 18 episodes of candidemia. J Infect. 2016;1–24.

7. Sharma C, Kumar N, Pandey R, Meis JF, Chowdhary A. Whole genome sequencing of emerging multidrug resistant Candida auris isolates in India demonstrates low genetic variation. New Microbes New Infect. 2016;13(C):77–82.

8. Belkin A, Gazit Z, Keller N, Ben-Ami R, Wieder-Finesod A, Novikov A, et al. Candida auris Infection Leading to Nosocomial Transmission, Israel, 2017. Emerg Infect Dis J. 2018;24(4):801–4.

9. Schelenz S, Hagen F, Rhodes JL, Abdolrasouli A, Chowdhary A, Hall A, et al. First hospital outbreak of the globally emerging Candida auris in a European hospital. Antimicrob Resist Infect Control. 2016;1–7.

10. Ruiz-Gaitan A, Moret AM, Tasias-Pitarch M, López AIA, Martínez-Morel H, Calabuig E, et al. An outbreak due to Candida auris with prolonged colonization and candidemia in a tertiary care European hospital. Mycoses. 2018;1–19.

11. Escandon P, Chow NA, Caceres DH, Gade L, Berkow EL, Armstrong P, et al. Molecular Epidemiology of Candida auris in Colombia Reveals a Highly Related, Countrywide Colonization With Regional Patterns in Amphotericin B Resistance. Clin Infect Dis. 2018;13:e1006290–7.

12. Welsh RM, Bentz ML, Shams A, Houston H, Lyons A, Rose LJ, et al. Survival, Persistence, and Isolation of the Emerging Multidrug-Resistant Pathogenic Yeast Candida auris on a Plastic Health Care Surface. J Clin Microbiol. 2017;55(10):2996–3005.

13. Lee WG, Shin JH, Uh Y, Kang MG, Kim SH, Park KH, et al. First three reported cases of nosocomial fungemia caused by Candida auris. J Clin Microbiol. 2011;49(9):3139–42.

14. Chow NA, Gade L, Tsay SV, Forsberg K, Greenko JA, Southwick KL, et al. Multiple introductions and subsequent transmission of multidrug-resistant Candida auris in the USA: a molecular epidemiological survey. Lancet Infect Dis. 2018 Dec 1;18(12):1377–84.

15. Chow NA, Groot T de, Badali H, Abastabar M, Chiller TM, Meis JF. Potential Fifth Clade of Candida auris, Iran, 2018 - Volume 25, Number 9—September 2019 - Emerging Infectious Diseases journal - CDC. [cited 2022 Aug 17]; Available from: https://wwwnc.cdc.gov/eid/article/25/9/19-0686_article

16. Muñoz JF, Gade L, Chow NA, Loparev VN, Juieng P, Berkow EL, et al. Genomic insights into multidrug-resistance, mating and virulence in Candida auris and related emerging species. Nat Commun. 2018;9(1):5346.

17. Chow NA, Muñoz JF, Gade L, Berkow EL, Li X, Welsh RM, et al. Tracing the Evolutionary History and Global Expansion of Candida auris Using Population Genomic Analyses. mBio. 2020;11(2):e03364–19.

18. Chowdhary A, Prakash A, Sharma C, Kordalewska M, Kumar A, Sarma S, et al. A multicentre study of antifungal susceptibility patterns among 350 Candida auris isolates (2009–17) in India: role of the ERG11 and FKS1 genes in azole and echinocandin resistance. J Antimicrob Chemother. 2018;13:e1006290.-9.

19. Rhodes J, Abdolrasouli A, Farrer RA, Cuomo CA, Aanensen DM, Armstrong-James D, et al. Genomic epidemiology of the UK outbreak of the emerging human fungal pathogen Candida auris. Emerg Microbes Infect. 2018;7(1):43.

20. Berkow EL, Lockhart SR. Activity of CD101, a long-acting echinocandin, against clinical isolates of Candida auris. Diagn Microbiol Infect Dis. 2017;

21. Kathuria S, Singh PK, Sharma C, Prakash A, Masih A, Kumar A, et al. Multidrug-Resistant Candida auris Misidentified as Candida haemulonii: Characterization by Matrix-Assisted Laser Desorption Ionization-Time of Flight Mass Spectrometry and DNA Sequencing and Its Antifungal Susceptibility Profile Variability by Vitek 2, CLSI Broth Microdilution, and Etest Method. J Clin Microbiol. 2015;53(6):1823–30.

22. Chowdhary A, Kumar VA, Sharma C, Prakash A, Agarwal K, Babu R, et al. Multidrug-resistant endemic clonal strain of Candida auris in India. Eur J Clin Microbiol Infect Dis Off Publ Eur Soc Clin Microbiol. 2013;33(6):919–26.

23. Centre for Disease Control E. Candida auris in healthcare settings – Europe. 2018;1– 10.

24. CDC. Centers for Disease Control and Prevention. 2022 [cited 2022 Aug 17]. The biggest antibiotic-resistant threats in the U.S. Available from: https://www.cdc.gov/drugresistance/biggest-threats.html

25. Taori SK, Khonyongwa K, Hayden I, Athukorala GDA, Letters A, Fife A, et al. Candida auris outbreak: Mortality, interventions and cost of sustaining control. J Infect. 2019;1–26.

26. Borman AM, Johnson EM. Candida auris in the UK: Introduction, dissemination, and control. PLOS Pathog. 2020 Jul 30;16(7):e1008563.

27. Borman AM, Szekely A, Johnson EM. Isolates of the emerging pathogen Candida auris present in the UK have several geographic origins. Med Mycol. 2017;55(5):53–567.

28. Pacilli M, Kerins JL, Clegg WJ, Walblay KA, Adil H, Kemble SK, et al. Regional Emergence of Candida auris in Chicago and Lessons Learned From Intensive Follow-up at 1 Ventilator-Capable Skilled Nursing Facility. Clin Infect Dis. 2020;71(11):e718–725.

29. Rex JH, Alexander BD, Andes D, Arthington-Skaggs B. Reference method for broth dilution antifungal susceptibility testing of yeasts; Approved standard—third edition (M27-A3). 2008. (Clinical and Laboratory Standards Institute (CLSI)).

30. Arendrup MC, Guinea J, Cuenca-Estrella M, Meletiadis J, Mouton JW, Lagrou K, et al. EUCAST DEFINITIVE DOCUMENT E.DEF 9.3. 2015;23.

31. Li H, Durbin R. Fast and accurate short read alignment with Burrows-Wheeler transform. Bioinformatics. 2009;25(14):1754–60.

32. Li H, Handsaker B, Wysoker A, Fennell T, Ruan J, Homer N, et al. The Sequence Alignment/Map format and SAMtools. Bioinformatics. 2009;25(16):2078–9.

33. Picard [Internet]. Available from: http://broadinstitute.github.io/picard

34. McKenna A, Hanna M, Banks E, Sivachenko A, Cibulskis K, Kernytsky A, et al. The Genome Analysis Toolkit: A MapReduce framework for analyzing next-generation DNA sequencing data. Genome Res. 2010;20(9):1297–303.

35. Stamatakis A. RAxML-VI-HPC: maximum likelihood-based phylogenetic analyses with thousands of taxa and mixed models. Bioinformatics. 2006;22(21):2688–90.

36. Cingolani P, Platts A, Wang LL, Coon M, Nguyen T, Wang L, et al. A program for annotating and predicting the effects of single nucleotide polymorphisms, SnpEff: SNPs in the genome of Drosophila melanogaster strain w 1118D; iso-2; iso-3. Fly (Austin). 2012 Apr;6(2):80–92.

37. Rambaut A, Lam TT, Carvalho LM, Pybus OG. Exploring the temporal structure of heterochronous sequences using TempEst (formerly Path-O-Gen). Virus Evol. 2016;2(1):vew007-7.

38. Didelot X, Fraser C, Gardy J, Colijn C. Genomic infectious disease epidemiology in partially sampled and ongoing outbreaks. 2016 p. 1–21.

39. Plummer M, Best N, Cowles K, Vines K. CODA: convergence diagnosis and output analysis for MCMC. R News. 2006 Mar;6(1):7–11.

40. Calo S, Shertz-Wall C, Lee SC, Bastidas RJ, Nicolás FE, Granek JA, et al. Antifungal drug resistance evoked via RNAi-dependent epimutations. Nature. 2014;513(7519):555–8.

41. Chang Z, Billmyre RB, Lee SC, Heitman J. Broad antifungal resistance mediated by RNAi-dependent epimutation in the basal human fungal pathogen Mucor circinelloides. Chowdhary A, editor. PLOS Genet. 2019 Feb 11;15(2):e1007957.

42. Edwards HM, Rhodes J. Accounting for the Biological Complexity of Pathogenic Fungi in Phylogenetic Dating. J Fungi. 2021 Aug 14;7(8):661.

43. Seo K, Akiyoshi H, Ohnishi Y. Alteration of Cell Wall Composition Leads to Amphotericin B Resistance in Aspergillus flavus. Microbiol Immunol. 1999;43(11):1017– 25.

44. Didelot X, Kendall M, Xu Y, White PJ, McCarthy N. Genomic Epidemiology Analysis of Infectious Disease Outbreaks Using TransPhylo. Curr Protoc [Internet]. 2021 Feb [cited 2023 Jun 1];1(2). Available from: https://onlinelibrary.wiley.com/doi/10.1002/cpz1.60

