## Supplementary material for "Genomic epidemiology of *Candida auris* introduction and outbreaks in the United Kingdom": SuppInfo

**Supplementary Figure 1 – Regression plots a)** Clade I and **b)** Clade III

**Supplementary Figure 2 – Maximum likelihood phylogeny of all 207 isolates included within this study**

**Supplementary Figure 3 - Matrices of probability of direct transmission for all pairs of individual isolates within Clade I (a) and III (b).** These matrices show the probability of direct transmission between isolates with no probability in dark pink and 100% probability in turquoise. Sample names were removed**.** Tables 3 (Clade I) and 4 (Clade III) in the appendix include information on transmission patterns including probability, collection dates, hospitals, sources and patient numbers.

**Supplementary Figure 4 – Maximum likelihood phylogeny of 24 isolates from 4 London hospitals, including sequential isolates from single patients.** Bootstrap iterations over 1000 replicates performed on WGS SNPs data, where branch lengths represent average number of SNPs. Bootstrap support below 75% indicated on branches.

**Supplementary Table 1 – *C. auris* isolates included in this study, summarised by Clade and hospital centre, with project accession number for data repository. *ERG11* mutations and minimum inhibitory concentrations (MICs) also provided: FLC=fluconazole; ITR=itraconazole; VRC=voriconazole; POS=posaconazole; CAS=caspofungin; AMB=amphotericin B; FLY=5-flucytosine; blank=not done/available.**

| **Clade** | **Hospital Centre** | **Isolate** | **Sample Collection Date** | ***ERG11* mutation** | **FLC** | **ITR** | **VRC** | **POS** | **CAS** | **AMB** | **FLY** | **Discriminant Analysis of Principal Components (DAPC) Cluster Number** |
| --- | --- | --- | --- | --- | --- | --- | --- | --- | --- | --- | --- | --- |
| I | Chelsea & Westminster (PRJEB36822) | CA_ICA1 | 26/07/2016 | Y132F | 16 | <0.03 | 0.03 | 0.008 | 0.125 | 0.5 | <0.06 | 1 |
|  | Hammersmith (PRJEB36822) | CA_IC11  CA_IC12  CA_IC13  CA_IC14  CA_IC15  CA_IC16  CA_IC17  CA_IC18  CA_IC19  CA_IC20  CA_IC21  CA_IC22  CA_IC23 | 14/12/2018  09/12/2018  14/12/2018  22/11/2018  30/12/2018  30/12/2018  04/01/2019  01/02/2019  05/02/2019  04/02/2019  05/02/2019  19/02/2019  19/02/2019 | K143R  K143R  K143R  K143R  K143R  K143R  K143R  K143R  K143R  K143R  K143R  K143R  K143R | 128  >64  128  128  64  128  8  >64  8 | 0.06  0.125  0.125  0.06  0.06  0.06  <0.03  0.25  <0.03 | 0.125  1  0.125  0.125  0.125  0.125  <0.008  1  0.015 | 0.03  0.125  0.06  0.03  0.06  0.03  <0.008  0.25  <0.008 | 0.06  0.5  0.125  0.06  0.06  0.125  0.125  0.5  0.125 | 0.5  0.5  1  0.5  0.5  0.5  2  1  1 | <0.06  0.25  <0.06  <0.06  <0.06  <0.06  <0.06  0.25  <0.06 | 1  1  1  1  1  1  1  1  3  1  1  1  1 |
|  | King’s College (PRJEB36563) | PHE_101  PHE_33  PHE_64  PHE_66  PHE_70  PHE_80  PHE_81  PHE_87  PHE_88  PHE_91  PHE_92  PHE_93  PHE_97  PHE_98  PHE_99  Sample_10  Sample_11  Sample_12  Sample_13  Sample_14  Sample_17  Sample_18  Sample_2  Sample_20  Sample_21  Sample_22  Sample_23  Sample_3  Sample_30  Sample_35  Sample_36  Sample_37  Sample_38  Sample_4  Sample_41  Sample_42  Sample_43  Sample_44  Sample_45  Sample_46  Sample_47  Sample_48  Sample_49  Sample_5  Sample_50  Sample_52  Sample_53  Sample_55  Sample_8 | 25/10/2016  28/07/2016  29/08/2016  08/09/2016  14/09/2016  27/09/2016  03/10/2016  16/10/2016  19/10/2016  30/10/2016  16/10/2016  20/10/2016  17/10/2016  24/10/2016  14/10/2016  09/01/2017  06/01/2018  21/09/2017  28/12/2016  14/09/2016  18/07/2018  20/07/2018  15/10/2016  17/10/2016  14/10/2016  22/06/2018  07/06/2018  24/10/2016  25/05/2018  30/05/2018  14/06/2018  18/06/2018  16/06/2018  06/10/2016  27/05/2018  15/05/2018  27/05/2018  01/06/2018  26/05/2018  28/05/2018  31/05/2018  29/05/2018  20/05/2018  22/06/2017  21/05/2018  03/07/2018  03/07/2018  03/07/2018  01/11/2016 | Y132F  Y132F  Y132F  Y132F  Y132F  Y132F  Y132F  Y132F  Y132F  Y132F  Y132F  Y132F  Y132F  Y132F  Y132F  Y132F  Y132F  Y132F  Y132F  Y132F  Y132F  Y132F  Y132F  Y132F  Y132F  Y132F  Y132F  Y132F  Y132F  Y132F  Y132F  Y132F  Y132F  Y132F  Y132F  Y132F  Y132F  Y132F  Y132F  Y132F  Y132F  Y132F  Y132F  Y132F  Y132F  Y132F  Y132F  Y132F  Y132F | 16  32  64  >64  64  32  64  32  32  32  >64  64  >64  32  >64  32  >64  32  32  32  32  32  64 | 0.25  <0.03  0.06  0.03  0.03 | 0.125  0.125  0.5  0.5  0.25  0.125  0.5  0.5  0.25  0.125  0.25  0.5  0.25  0.125  2  0.5  >16  1  0.125  0.25  0.25  0.25  0.25 | 0.06  <0.03  0.06  0.03  0.03 | 0.125  1.5  >32  0.125  0.25  0.125  0.5  0.125  0.19  1  0.5  0.5  0.25  0.25  0.25  0.25  0.25  0.5  0.25 | 2  2  1  1  1  0.25  1  1  1  1  1  1  1  2  2  1  1  1  1  1  1  1  1  1  1  1  1  1  1  1  1 | <0.125  8  4  >64  4  1  >64  32  0.25  0.125  32  16  4  0.25  8  8  0.125  0.125  0.125  0.5  0.125  0.25  0.25  0.125  0.125  0.25 | 3  3  3  3  3  3  3  3  3  3  3  3  3  3  3  2  3  6  5  5  3  3  5  3  3  3  3  3  3  5  5  5  3  6  3  5  5  5  5  5  5  5  5  2  5  5  5  5  3 |
|  | Northwick Park (PRJEB36822) | CA_IC10  CA_IC24  CA_IC7  CA_IC8  CA_IC9 | 05/12/2017  08/05/2019  26/10/2017  26/10/2017  17/11/2017 | Y132F  Y132F  Y132F  Y132F  Y132F | 16  16  16  32 | <0.03  <0.03  <0.03  <0.03 | 0.03  0.03  0.125  0.03 | <0.008  <0.008  <0.03  <0.008 | 0.06  0.125  0.5  0.125 | 0.25  0.5  1  0.5 | <0.06  <0.06  <0.125  <0.06 | 3  1  1  1  1 |
|  | Royal Brompton (PRJEB26393, with PHE_ isolates submitted under PRJEB36822) | 15B6  15B10  15B5  16B27a  16B13  16B12  16B16  16B30  16B31  16B18  16B15a  16B20  16B21  16B22a  16B24b  16B22b  16B15b  16B24a  16B25  16B26  16I30  16I29a  16I29b  16I33  16I34  16I17  16I27b  PHE_20  PHE_23  PHE_24  PHE_25  PHE_28  PHE_34  PHE_35  PHE_50 | 22/10/2015  28/12/2015  19/10/2015  15/03/2016  11/01/2016  04/01/2016  21/02/2016  17/10/2016  16/10/2016  01/02/2016  06/02/2016  16/02/2016  22/02/2016  27/02/2016  07/03/2016  09/03/2016  09/03/2016  12/03/2016  13/03/2016  14/03/2016  21/03/2016  18/01/2016  18/01/2016  16/06/2016  23/10/2016  08/02/2016  14/03/2016  08/07/2016  08/07/2016  08/07/2016  08/07/2016  09/03/2016  26/07/2016  28/07/2016  14/04/2016 | Y132F  Y132F  Y132F  Y132F  Y132F  Y132F  Y132F  Y132F  Y132F  Y132F  Y132F  Y132F  Y132F  Y132F  Y132F  Y132F  Y132F  Y132F  Y132F  Y132F  Y132F  Y132F  Y132F  Y132F  Y132F  Y132F  Y132F  Y132F  Y132F  Y132F  Y132F  Y132F  Y132F  Y132F  Y132F | 256  256  256  256  256  256  256  128  256  256  16  256  16  256  8  256  64  256  256  128  256  >64  16  8 | 0.06  16  16  16  0.06  16  0.03  0.06  0.06  0.12  0.015  16  0.03  0.03  2  2  0.03  2  0.03  16  16  0.03  0.03  16  <0.03  0.03 | >16  0.125  0.25 | >16  <0.03  0.03 |  | 1  1  1 | >64  <0.125  >64  <0.125 | 5  3  3  5  5  5  4  3  3  5  5  3  5  3  3  3  2  5  3  3  3  3  5  3  3  5  4  3  3  3  2  2  3  3  3 |
|  | Stoke Mandeville (PRJEB36822) | PHE_26  PHE_27 | 14/12/2015  15/02/2016 | Y132F  Y132F | 32  16 | <0.03  <0.03 | 0.06  0.06 | <0.03  <0.03 |  | 2  1 | <0.125  <0.125 | 3  3 |

|  | West Middlesex  (PRJEB36822) | CA_IC2  CA_IC3  CA_IC4  CA_IC5  CA_IC6 | 03/12/2016  28/12/2016  03/01/2017  19/04/2019  17/05/2019 | Y132F  Y132F  Y132F  Y132F  Y132F | 64  64 | <0.03  <0.03 | 0.125  0.125 | <0.008  <0.008 | 0.125  0.125 | 1  1 | <0.06  <0.06 | 1  1  1  1  1 |
| --- | --- | --- | --- | --- | --- | --- | --- | --- | --- | --- | --- | --- |
|  | Wexham Park  (PRJEB36822) | PHE_12  PHE_19  PHE_55  PHE_82 | 11/01/2016  24/06/2016  22/08/2016  05/10/2016 | Y132F  Y132F  Y132F  Y132F | 16  16  16 | <0.03  <0.03  0.06 | 0.125  0.125  0.5 | <0.03  <0.03 | 0.06 | 1  1  0.5 | <0.125  <0.125 | 3  3  3  3 |
| III | John Radcliffe (Project Accession PRJ415955, with PHE_ isolates submitted under PRJEB36822) | PHE_104  PHE_111  PHE_2  PHE_32  PHE_8  PHE_84  PHE_85  PHE_86  PHE_95  SRR7976540  SRR7976541  SRR7976542  SRR7976543  SRR7976544  SRR7976545  SRR7976546  SRR7976547  SRR7976548  SRR7976549  SRR7976550  SRR7976551  SRR7976552  SRR7976553  SRR7976554  SRR7976555  SRR7976556  SRR7976557  SRR7976558  SRR7976559  SRR7976560  SRR7976561  SRR7976562  SRR7976563  SRR7976564  SRR7976565  SRR7976566  SRR7976567  SRR7976568  SRR7976569  SRR7976570  SRR7976571  SRR7976572  SRR7976573  SRR7976574  SRR7976575  SRR7976576  SRR7976577  SRR7976578  SRR7976579  SRR7976580  SRR7976581  SRR7976582  SRR7976583  SRR7976584  SRR7976585  SRR7976586  SRR7976587  SRR7976588  SRR7976589  SRR7976590  SRR7976591  SRR7976592  SRR7976593  SRR7976594  SRR7976595  SRR7976596  SRR7976597  SRR7976598  SRR7976599  SRR7976600  SRR7976601  SRR7976602  SRR7976603  SRR7976604  SRR7976605  SRR7976606  SRR7976607  SRR7976608  SRR7976609  SRR7976610  SRR7976611  SRR7976612  SRR7976613  SRR7976614  SRR7976615  SRR7976616  SRR7976617 | 25/10/2016  02/11/2016  27/05/2015  22/04/2016  02/02/2015  16/10/2016  29/10/2015  13/07/2016  16/10/2016  10/02/2017  05/05/2017  17/11/2016  15/01/2017  03/04/2017  25/01/2017  13/02/2017  30/01/2017  30/08/2017  02/02/2015  19/12/2016  20/02/2017  17/02/2017  18/03/2017  10/02/2017  30/01/2017  22/05/2017  25/10/2016  07/04/2017  22/03/2017  16/05/2017  10/05/2017  14/04/2017  16/01/2017  10/01/2017  27/03/2017  24/01/2017  31/05/2017  30/01/2017  15/02/2017  29/01/2017  16/05/2017  10/03/2017  15/01/2017  09/01/2017  05/04/2017  07/05/2017  19/12/2016  27/01/2017  10/07/2015  26/04/2017  20/02/2017  24/11/2015  04/04/2017  28/11/2016  03/04/2017  15/01/2017  13/02/2017  08/03/2017  17/02/2017  15/03/2017  11/12/2016  24/03/2017  04/04/2017  30/01/2017  17/07/2017  17/05/2017  05/12/2016  09/04/2017  25/01/2017  29/03/2017  16/05/2017  16/05/2017  24/04/2017  09/01/2017  08/02/2017  30/01/2017  10/04/2017  13/02/2017  29/01/2017  10/04/2017  13/02/2017  13/02/2017  25/01/2017  02/01/2016  31/03/2017  13/07/2015  22/03/2017 | F126L  F126L  F126L  F126L  F126L  F126L  F126L  F126L  F126L  F126L  F126L  F126L  F126L  F126L  F126L  F126L  F126L  F126L  F126L  F126L  F126L  F126L  F126L  F126L  F126L  F126L  F126L  F126L  F126L  F126L  F126L  F126L  F126L  F126L  F126L  F126L  F126L  F126L  F126L  F126L  F126L  F126L  F126L  F126L  F126L  F126L  F126L  F126L  F126L  F126L  F126L  F126L  F126L  F126L  F126L  F126L  F126L  F126L  F126L  F126L  F126L  F126L  F126L  F126L  F126L  F126L  F126L  F126L  F126L  F126L  F126L  F126L  F126L  F126L  F126L  F126L  F126L  F126L  F126L  F126L  F126L  F126L  F126L  F126L  F126L  F126L  F126L | >64  >64  >64  >64  >64  >64  >64  >64  >64 | 0.5  0.5  0.25  0.25  0.25  0.25  0.5  0.25  0.5 | 4  8  1  4  2  2  4  2  4 | 0.25  0.125  0.06  0.06  0.06  0.06  0.06  0.06  0.06 |  | 1  1  4  1  1  1  2  1  2 | 0.25  0.25  <0.125  0.5  <0.125  1  <0.125  <0.125  0.25 | 1  1  1  1  1  1  1  1  1  5  1  1  5  2  5  1  5  5  5  1  5  1  5  5  5  5  1  3  5  1  1  5  6  5  5  5  1  5  6  1  5  5  1  5  5  5  2  1  1  2  1  1  5  5  5  5  1  3  1  3  1  4  1  5  4  5  1  5  5  5  4  1  1  1  5  1  1  1  1  5  5  5  2  1  5  1  5 |
|  | King’s College (Project Accession PRJEB36563) | Sample_16  Sample_24  Sample_57 | 18/07/2018  21/07/2018  04/07/2018 | F126L  F126L  F126L | 64  64  64 |  | 2  1  2 |  |  | 1  0.5  1 |  | 1  1  1 |
|  | Wexham Park (PRJEB36822) | PHE_63  PHE_89 | 02/09/2016  16/05/2016 | F126L  F126L | >64 |  | 4 |  | 0.25 | 0.25 |  | 1  1 |
|  | HCA laboratories (PRJEB36822) | PHE_6 | 10/06/2014 | F126L | >64 | 1 | 4 | 0.125 |  | 1 | <0.125 | 1 |

**Supplementary Table 2 – Average single nucleotide polymorphism (SNP) distance between isolates on average in individual hospital centres and within each clade. Hospital centres with no isolates for a particular clade are denoted as ‘-’.**

|  | **Clade Total** | **Chelsea and Westminster** | **Hammersmith** | **John Radcliffe** | **King’s College** | **Northwick Park** | **Royal Brompton** | **Stoke Mandeville** | **West Middlesex** | **Wexham Park** |
| --- | --- | --- | --- | --- | --- | --- | --- | --- | --- | --- |
| Clade I | 139 | Only 1 isolate in hospital | 14 | - | 119 | 7 | 63 | 14 | 8 | 69 |
| Clade III | 151 | - | - | 166 | 1 | - | - | - | - | 22 |

| **Supplementary Table 3: Transmission between Clade I isolates using *TransPhylo* with a probability threshold above 75%.** The transmission pathway is from isolate 1 to isolate 2 including probability of transmission from isolates, date of isolate collection, hospital, source and patient numbers. Patient numbers correspond to what was provided by each hospital centre. | | | | | | | | | | | |
| --- | --- | --- | --- | --- | --- | --- | --- | --- | --- | --- | --- |
| Transmission --> | |  |  |  |  |  |  |  |  |  |  |
| Isolate 1 | Isolate 2 | Probability of Transmission from Isolate 1 to Isolate 2 | Date of Collection Isolate 1 | Date of Collection Isolate 2 | Hospital Isolate 1 | Hospital Isolate 2 | Source Isolate 1 | Source Isolate 2 | Patient Number Isolate 1 | Patient Number Isolate 2 | Notes |
| 16B9044 | 16B9260 | 1 | 07/03/2016 | 09/03/2016 | Royal Brompton | Royal Brompton | Clinical | Clinical | Patient C | Patient A |  |
| 16I10532 | 16B6373 | 1 | 21/03/2016 | 16/02/2016 | Royal Brompton | Royal Brompton | Clinical | Clinical | Patient B | Unknown | Collection Date of Isolate 2 before Isolate 1 |
| 16I10532 | 16B7775 | 1 | 21/03/2016 | 27/02/2016 | Royal Brompton | Royal Brompton | Clinical | Clinical | Patient B | Patient A | Collection Date of Isolate 2 before Isolate 1 |
| CA_IC7 | CA_IC8 | 1 | 26/10/2017 | 26/10/2017 | Northwick Park | Northwick Park | Clinical | Clinical | 3 | 3 | Same collection date, same patient |
| PHE_19 | PHE_50 | 1 | 24/06/2016 | 14/04/2016 | Wexham Park | Royal Brompton | Clinical | Clinical | Unknown | Unknown | Collection Date of Isolate 2 before Isolate 1 |
| Sample_13 | Sample_53 | 1 | 28/12/2016 | 03/07/2018 | King’s College | King’s College | Clinical | Environment | 24 | N/A |  |
| Sample_36 | Sample_45 | 1 | 14/06/2018 | 26/05/2018 | King’s College | King’s College | Environment | Clinical | N/A | 39 | Collection Date of Isolate 2 before Isolate 1 |
| Sample_42 | Sample_46 | 1 | 15/05/2018 | 28/05/2018 | King’s College | King’s College | Clinical | Clinical | 37 | 42 |  |
| Sample_48 | Sample_44 | 1 | 29/05/2018 | 01/06/2018 | King’s College | King’s College | Clinical | Clinical | 40 | 45 |  |
| Sample_50 | Sample_43 | 1 | 21/05/2018 | 27/05/2018 | King’s College | King’s College | Clinical | Clinical | 31 | 38 |  |
| Sample_50 | Sample_42 | 1 | 21/05/2018 | 15/05/2018 | King’s College | King’s College | Clinical | Clinical | 31 | 37 | Collection Date of Isolate 2 before Isolate 1 |
| Sample_50 | Sample_47 | 1 | 21/05/2018 | 31/05/2018 | King’s College | King’s College | Clinical | Clinical | 31 | 43 |  |
| PHE_99 | Sample_21 | 0.98120376 | 14/10/2016 | 14/10/2016 | King’s College | King’s College | Clinical | Clinical | 17 | 8 |  |
| PHE_23 | PHE_101 | 0.93881224 | 08/07/2016 | 25/10/2016 | Royal Brompton | King’s College | Clinical | Clinical | Unknown | 16 |  |
| CA_IC24 | CA_IC5 | 0.93661268 | 08/05/2019 | 19/04/2019 | Northwick Park | West Middlesex | Clinical | Clinical | 3 | 2 | Collection Date of Isolate 2 before Isolate 1 |
| CA_IC24 | CA_IC9 | 0.9330134 | 08/05/2019 | 17/11/2017 | Northwick Park | Northwick Park | Clinical | Clinical | 3 | 3 | Collection Date of Isolate 2 before Isolate 1 |
| CA_IC6 | CA_IC4 | 0.93181364 | 17/05/2019 | 03/01/2017 | West Middlesex | West Middlesex | Clinical | Clinical | 2 | 2 | Collection Date of Isolate 2 before Isolate 1 |
| PHE_81 | PHE_19 | 0.90541892 | 03/10/2016 | 24/06/2016 | King’s College | Wexham Park | Clinical | Clinical | 7 | Unknown | Collection Date of Isolate 2 before Isolate 1 |
| PHE_81 | 16I33 | 0.90541892 | 03/10/2016 | 16/06/2016 | King’s College | Royal Brompton | Clinical | Unknown | 7 | Unknown | Collection Date of Isolate 2 before Isolate 1 |
| PHE_81 | PHE_98 | 0.90541892 | 03/10/2016 | 24/10/2016 | King’s College | King’s College | Clinical | Clinical | 7 | 15 |  |
| PHE_81 | PHE_12 | 0.90241952 | 03/10/2016 | 11/01/2016 | King’s College | Wexham Park | Clinical | Unknown | 7 | Unknown | Collection Date of Isolate 2 before Isolate 1 |
| Sample_2 | CA_IC10 | 0.89982004 | 15/10/2016 | 05/12/2017 | King’s College | Northwick Park | Clinical | Clinical | 7 | 3 |  |
| 16B31 | PHE_24 | 0.889822 | 16/10/2016 | 08/07/2016 | Royal Brompton | Royal Brompton | Unknown | Clinical | Unknown | Unknown | Collection Date of Isolate 2 before Isolate 1 |
| Sample_8 | Sample_22 | 0.8822236 | 01/11/2016 | 22/06/2018 | King’s College | King’s College | Clinical | Clinical | 5 | 50 |  |
| PHE_28 | PHE_25 | 0.86382723 | 09/03/2016 | 08/07/2016 | Royal Brompton | Royal Brompton | Clinical | Clinical | Unknown | Unknown |  |
| 16B4119 | 16B10136 | 0.8634273 | 01/02/2016 | 15/03/2016 | Royal Brompton | Royal Brompton | Clinical | Clinical | Unknown | Unknown |  |
| 16B4119 | 16I5822 | 0.8634273 | 01/02/2016 | 08/02/2016 | Royal Brompton | Royal Brompton | Clinical | Clinical | Unknown | Unknown |  |
| Sample_37 | Sample_55 | 0.8620276 | 18/06/2018 | 03/07/2018 | King’s College | King’s College | Clinical | Environment | 49 | N/A |  |
| PHE_81 | 16B9044 | 0.85342931 | 03/10/2016 | 07/03/2016 | King’s College | Royal Brompton | Clinical | Clinical | 7 | Patient C | Collection Date of Isolate 2 before Isolate 1 |
| PHE_64 | PHE_66 | 0.8362328 | 29/08/2016 | 08/09/2016 | King’s College | King’s College | Clinical | Clinical | 4 | 5 |  |
| PHE_64 | PHE_55 | 0.83623275 | 29/08/2016 | 22/08/2016 | King’s College | Wexham Park | Clinical | Clinical | 4 | Unknown | Collection Date of Isolate 2 before Isolate 1 |
| PHE_35 | PHE_33 | 0.835233 | 28/07/2016 | 28/07/2016 | Royal Brompton | King’s College | Clinical | Clinical | Unknown | 3 |  |
| CA_IC13 | CA_IC11 | 0.83263347 | 14/12/2018 | 14/12/2018 | Hammersmith | Hammersmith | Clinical | Clinical | 4 | 4 | Same collection date, same patient |
| Sample_12 | Sample_4 | 0.82483503 | 21/09/2017 | 06/10/2016 | King’s College | King’s College | Clinical | Clinical | 34 | 7 |  |
| Sample_10 | PHE_82 | 0.7988402 | 09/01/2017 | 05/10/2016 | King’s College | Wexham Park | Clinical | Clinical | 25 | Unknown | Collection Date of Isolate 2 before Isolate 1 |
| Sample_10 | Sample_5 | 0.7988402 | 09/01/2017 | 22/06/2017 | King’s College | King’s College | Clinical | Clinical | 25 | 7 |  |
| PHE_91 | PHE_87 | 0.790042 | 30/10/2016 | 16/10/2016 | King’s College | King’s College | Clinical | Clinical | 8 | 10 | Collection Date of Isolate 2 before Isolate 1 |
| PHE_81 | PHE_35 | 0.77864427 | 03/10/2016 | 28/07/2016 | King’s College | Royal Brompton | Clinical | Clinical | 7 | Unknown | Collection Date of Isolate 2 before Isolate 1 |
| CA_IC2 | CA_IC6 | 0.77704459 | 03/12/2016 | 17/05/2019 | West Middlesex | West Middlesex | Clinical | Clinical | 2 | 2 | Same patient |
| CA_IC13 | CA_IC18 | 0.7554489 | 14/12/2018 | 01/02/2019 | Hammersmith | Hammersmith | Clinical | Clinical | 4 | 5 |  |

| **Supplementary Table 4: Transmission between Clade III isolates using *TransPhylo* with a probability threshold above 75%.** The transmission pathway is from isolate 1 to isolate 2 including probability of transmission from isolates, date of isolate collection, hospital, source and patient numbers. Patient numbers correspond to what was provided by each hospital centre. | | | | | | | | | | | |
| --- | --- | --- | --- | --- | --- | --- | --- | --- | --- | --- | --- |
| Transmission --> | |  |  |  |  |  |  |  |  |  |  |
| Isolate 1 | Isolate 2 | Probability of Transmission from Isolate 1 to Isolate 2 | Date of Collection Isolate 1 | Date of Collection Isolate 2 | Hospital Isolate 1 | Hospital Isolate 2 | Source Isolate 1 | Source Isolate 2 | Patient Number Isolate 1 | Patient Number Isolate 2 | Notes |
| PHE_104 | PHE_86 | 0.8494301 | 25/10/2016 | 13/07/2016 | John Radcliffe | John Radcliffe | Clinical | Clinical | Unknown | Unknown | Collection Date of Isolate 2 before Isolate 1 |
| PHE_104 | PHE_95 | 0.8494301 | 25/10/2016 | 16/10/2016 | John Radcliffe | John Radcliffe | Clinical | Clinical | Unknown | Unknown | Collection Date of Isolate 2 before Isolate 1 |
| PHE_104 | SRR7976584 | 0.8494301 | 25/10/2016 | 28/11/2016 | John Radcliffe | John Radcliffe | Clinical | Clinical | Unknown | Patient_008 |  |
| PHE_6 | SRR7976616 | 0.95001 | 10/06/2014 | 13/07/2015 | Unknown | John Radcliffe | Clinical | Clinical | Unknown | Patient_003 |  |
| PHE_85 | SRR7976582 | 0.984803 | 29/10/2015 | 24/11/2015 | John Radcliffe | John Radcliffe | Clinical | Clinical | Unknown | Patient_004 |  |
| Sample_16 | Sample_24 | 0.8956209 | 18/07/2018 | 21/07/2018 | King’s College | King’s College | Clinical | Clinical | 52 | 54 |  |
| Sample_16 | Sample_57 | 0.8956209 | 18/07/2018 | 04/07/2018 | King’s College | King’s College | Clinical | Clinical | 52 | 21 | Collection Date of Isolate 2 before Isolate 1 |
| Sample_16 | SRR7976548 | 0.8956209 | 18/07/2018 | 30/08/2017 | King’s College | John Radcliffe | Clinical | Clinical | 52 | Patient_021 |  |
| Sample_16 | SRR7976571 | 0.8956209 | 18/07/2018 | 16/05/2017 | King’s College | John Radcliffe | Clinical | Environmental | 52 | N/A | Collection Date of Isolate 2 before Isolate 1 |
| SRR7976544 | SRR7976563 | 0.9998 | 03/04/2017 | 16/01/2017 | John Radcliffe | John Radcliffe | Clinical | Clinical | Patient_024 | Patient_014 | Collection Date of Isolate 2 before Isolate 1 |
| SRR7976546 | SRR7976552 | 0.77964407 | 13/02/2017 | 17/02/2017 | John Radcliffe | John Radcliffe | Clinical | Clinical | Patient_006 | Patient_019 |  |
| SRR7976546 | SRR7976570 | 0.7796441 | 13/02/2017 | 29/01/2017 | John Radcliffe | John Radcliffe | Clinical | Clinical | Patient_006 | Patient_009 | Collection Date of Isolate 2 before Isolate 1 |
| SRR7976554 | SRR7976551 | 1 | 10/02/2017 | 20/02/2017 | John Radcliffe | John Radcliffe | Clinical | Clinical | Patient_018 | Patient_020 |  |
| SRR7976554 | SRR7976564 | 0.9876025 | 10/02/2017 | 10/01/2017 | John Radcliffe | John Radcliffe | Clinical | Clinical | Patient_018 | Patient_013 | Collection Date of Isolate 2 before Isolate 1 |
| SRR7976554 | SRR7976596 | 1 | 10/02/2017 | 17/05/2017 | John Radcliffe | John Radcliffe | Clinical | Clinical | Patient_018 | Patient_028 |  |
| SRR7976557 | PHE_32 | 1 | 25/10/2016 | 22/04/2016 | John Radcliffe | John Radcliffe | Clinical | Clinical | Patient_006 | Unknown | Collection Date of Isolate 2 before Isolate 1 |
| SRR7976557 | SRR7976542 | 1 | 25/10/2016 | 17/11/2016 | John Radcliffe | John Radcliffe | Clinical | Clinical | Patient_006 | Patient_007 |  |
| SRR7976557 | SRR7976573 | 1 | 25/10/2016 | 15/01/2017 | John Radcliffe | John Radcliffe | Clinical | Clinical | Patient_006 | Patient_006 |  |
| SRR7976558 | SRR7976572 | 0.8116377 | 07/04/2017 | 10/03/2017 | John Radcliffe | John Radcliffe | Clinical | Clinical | Patient_029 | Patient_022 | Collection Date of Isolate 2 before Isolate 1 |
| SRR7976562 | SRR7976575 | 0.8120376 | 14/04/2017 | 05/04/2017 | John Radcliffe | John Radcliffe | Clinical | Clinical | Patient_025 | Patient_025 | Collection Date of Isolate 2 before Isolate 1 |
| SRR7976562 | SRR7976583 | 0.8070386 | 14/04/2017 | 04/04/2017 | John Radcliffe | John Radcliffe | Clinical | Environmental | Patient_025 | N/A | Collection Date of Isolate 2 before Isolate 1 |
| SRR7976562 | SRR7976612 | 0.8120376 | 14/04/2017 | 13/02/2017 | John Radcliffe | John Radcliffe | Clinical | Clinical | Patient_025 | Patient_006 | Collection Date of Isolate 2 before Isolate 1 |
| SRR7976565 | SRR7976540 | 0.8546291 | 27/03/2017 | 10/02/2017 | John Radcliffe | John Radcliffe | Clinical | Clinical | Patient_025 | Patient_017 | Collection Date of Isolate 2 before Isolate 1 |
| SRR7976565 | SRR7976553 | 0.8546291 | 27/03/2017 | 18/03/2017 | John Radcliffe | John Radcliffe | Clinical | Clinical | Patient_025 | Patient_023 | Collection Date of Isolate 2 before Isolate 1 |
| SRR7976565 | SRR7976554 | 0.8546291 | 27/03/2017 | 10/02/2017 | John Radcliffe | John Radcliffe | Clinical | Clinical | Patient_025 | Patient_018 | Collection Date of Isolate 2 before Isolate 1 |
| SRR7976565 | SRR7976558 | 0.8116377 | 27/03/2017 | 07/04/2017 | John Radcliffe | John Radcliffe | Clinical | Clinical | Patient_025 | Patient_029 |  |
| SRR7976565 | SRR7976568 | 0.8478304 | 27/03/2017 | 30/01/2017 | John Radcliffe | John Radcliffe | Clinical | Clinical | Patient_025 | Patient_010 | Collection Date of Isolate 2 before Isolate 1 |
| SRR7976567 | SRR7976541 | 0.96780644 | 31/05/2017 | 05/05/2017 | John Radcliffe | John Radcliffe | Clinical | Clinical | Patient_022 | Patient_033 | Collection Date of Isolate 2 before Isolate 1 |
| SRR7976567 | SRR7976546 | 0.7796441 | 31/05/2017 | 13/02/2017 | John Radcliffe | John Radcliffe | Clinical | Clinical | Patient_022 | Patient_006 | Collection Date of Isolate 2 before Isolate 1 |
| SRR7976567 | SRR7976560 | 0.96780644 | 31/05/2017 | 16/05/2017 | John Radcliffe | John Radcliffe | Clinical | Environmental | Patient_022 | N/A | Collection Date of Isolate 2 before Isolate 1 |
| SRR7976567 | SRR7976561 | 0.96780644 | 31/05/2017 | 10/05/2017 | John Radcliffe | John Radcliffe | Clinical | Clinical | Patient_022 | Patient_033 | Collection Date of Isolate 2 before Isolate 1 |
| SRR7976567 | SRR7976603 | 0.96780644 | 31/05/2017 | 24/04/2017 | John Radcliffe | John Radcliffe | Clinical | Clinical | Patient_022 | Patient_037 | Collection Date of Isolate 2 before Isolate 1 |
| SRR7976567 | SRR7976609 | 0.94921016 | 31/05/2017 | 29/01/2017 | John Radcliffe | John Radcliffe | Clinical | Clinical | Patient_022 | Patient_009 |  |
| SRR7976576 | SRR7976610 | 1 | 07/05/2017 | 10/04/2017 | John Radcliffe | John Radcliffe | Clinical | Clinical | Patient_027 | Patient_029 |  |
| SRR7976576 | SRR7976615 | 1 | 07/05/2017 | 31/03/2017 | John Radcliffe | John Radcliffe | Clinical | Clinical | Patient_027 | Patient_027 | Collection Date of Isolate 2 before Isolate 1 |
| SRR7976588 | SRR7976590 | 0.75144971 | 08/03/2017 | 15/03/2017 | John Radcliffe | John Radcliffe | Clinical | Clinical | Patient_019 | Patient_021 |  |
| SRR7976593 | SRR7976589 | 1 | 04/04/2017 | 17/02/2017 | John Radcliffe | John Radcliffe | Environmental | Clinical | N/A | Patient_018 | Collection Date of Isolate 2 before Isolate 1 |
| SRR7976593 | SRR7976597 | 1 | 04/04/2017 | 05/12/2016 | John Radcliffe | John Radcliffe | Environmental | Clinical | N/A | Patient_009 | Collection Date of Isolate 2 before Isolate 1 |
| SRR7976608 | PHE_104 | 0.8494301 | 13/02/2017 | 25/10/2016 | John Radcliffe | John Radcliffe | Clinical | Clinical | Patient_010 | Unknown | Collection Date of Isolate 2 before Isolate 1 |
| SRR7976608 | PHE_111 | 0.9940012 | 13/02/2017 | 02/11/2016 | John Radcliffe | John Radcliffe | Clinical | Clinical | Patient_010 | Unknown | Collection Date of Isolate 2 before Isolate 1 |
| SRR7976608 | PHE_8 | 0.886228 | 13/02/2017 | 02/02/2015 | John Radcliffe | John Radcliffe | Clinical | Clinical | Patient_010 | Unknown | Collection Date of Isolate 2 before Isolate 1 |
| SRR7976608 | PHE_84 | 1 | 13/02/2017 | 16/10/2016 | John Radcliffe | John Radcliffe | Clinical | Clinical | Patient_010 | Unknown | Collection Date of Isolate 2 before Isolate 1 |
| SRR7976608 | SRR7976604 | 1 | 13/02/2017 | 09/01/2017 | John Radcliffe | John Radcliffe | Clinical | Clinical | Patient_010 | Patient_012 | Collection Date of Isolate 2 before Isolate 1 |
| SRR7976608 | SRR7976614 | 0.9566087 | 13/02/2017 | 02/01/2016 | John Radcliffe | John Radcliffe | Clinical | Clinical | Patient_010 | Patient_005 | Collection Date of Isolate 2 before Isolate 1 |
| SRR7976613 | SRR7976569 | 1 | 25/01/2017 | 15/02/2017 | John Radcliffe | John Radcliffe | Clinical | Clinical | Patient_011 | Patient_011 |  |

| **Supplementary Table 5: Sequential *C. auris* isolates (all Clade I) taken from patients within four London hospitals to investigate microevolution. MIC data can be found in Supp. Table 1.** | | | | |
| --- | --- | --- | --- | --- |
| **Hospital Centre** | **Patient** | **Isolate** | **Collection date** | **Sample site** |
| Chelsea & Westminster | 1 | CA_IC1 | 26/07/2016 | Groin swab |
| West Middlesex | 2 | CA_IC2 | 03/12/2016 | Urine |
|  |  | CA_IC3 | 28/12/2016 | Nose swab |
|  |  | CA_IC4 | 03/01/2017 | Nose swab |
|  |  | CA_IC5 | 19/04/2019 | Urine |
|  |  | CA_IC6 | 17/05/2019 | Urine |
| Northwick Park | 3 | CA_IC7 | 26/10/2017 | Exit site swab |
|  |  | CA_IC8 | 26/10/2017 | Throat swab |
|  |  | CA_IC9 | 17/11/2017 | Swab no site |
|  |  | CA_IC10 | 05/12/2017 | Exit site swab |
|  |  | CA_IC24 | 08/05/2019 | Exit site swab |
| Hammersmith | 4 | CA_IC11 | 14/12/2018 | Axilla swab |
|  |  | CA_IC12 | 09/12/2018 | Groin swab |
|  |  | CA_IC13 | 14/12/2018 | Groin swab |
|  | 5 | CA_IC14 | 22/11/2018 | Sputum |
|  |  | CA_IC15 | 30/12/2018 | Nose swab |
|  |  | CA_IC16 | 30/12/2018 | Groin swab |
|  |  | CA_IC17 | 04/01/2019 | Bronchial wash |
|  |  | CA_IC18 | 01/02/2019 | Urine |
|  |  | CA_IC20 | 04/02/2019 | Urine |
|  | 6 | CA_IC19 | 05/02/2019 | Rectal swab |
|  |  | CA_IC21 | 05/02/2019 | Groin swab |
|  |  | CA_IC22 | 19/02/2019 | Groin swab |
|  |  | CA_IC23 | 19/02/2019 | Urine |
